# Cultivation and characterization of a novel clade of deep-sea Chloroflexi: providing a glimpse of the phylum Chloroflexi involved in sulfur cycling

**DOI:** 10.1101/2021.01.05.425403

**Authors:** Rikuan Zheng, Ruining Cai, Rui Liu, Yeqi Shan, Ge Liu, Chaomin Sun

## Abstract

Chloroflexi bacteria are abundant and globally distributed in various unexplored biospheres on Earth. However, only few Chloroflexi members have been cultivated, hampering further understanding of this important group. In the current study, we firstly clarify the high abundance of the phylum Chloroflexi in deep-sea sediments via the operational taxonomic units analysis. We further successfully isolate a novel Chloroflexi strain ZRK33 from cold seep sediments by using an enrichment medium constantly supplemented with rifampicin. Phylogenetic analyses based on 16S rRNA gene, genome, RpoB and EF-tu proteins indicate that strain ZRK33 represents a novel class, and the class is designated as Sulfochloroflexia because whole set of genes encoding key enzymes responsible for assimilatory sulfate reduction are identified in the genome of strain ZRK33. Indeed, assimilation of sulfate or thiosulfate by strain ZRK33 evidently benefits its growth and morphogenesis. Proteomic results suggest that metabolization of sulfate or thiosulfate significantly promotes the transport and degradation of various macromolecules and thereby stimulating the energy production. Notably, the putative genes associated with assimilatory and dissimilatory sulfate reduction ubiquitously distribute in the metagenome-assembled genomes of 27 Chloroflexi members derived from deep-sea sediments, strongly suggesting that Chloroflexi bacteria play undocumented key roles in deep-sea sulfur cycling.

## Introduction

Deep marine subsurface is one of the least-understood habitats on Earth and is estimated to contain up to 3×10 ^29^ microbial cells, which is equivalent to the combined microbial biomass of the oceanic water column and terrestrial soil [1]. The prokaryotic biomass in deep marine subsurface sediments exceeds 10^5^ microbial cells/cm^3^ even at depths of nearly to 1,000 m below the seafloor [2,3]. These microorganisms are the primary drivers of elemental cycles within deep marine subsurface sediments and play key roles in the recycling of biogeochemical nutrients to the water column [4]. Members of the phylum Chloroflexi widely distributed in various environments with high abundance, for example, in some marine subsurface sediments the number of Chloroflexi bacteria is shown to be closely equivalent to other total bacterial counts [3,5–8], strongly suggesting that the phylum Chloroflexi is an essential group to maintain the population equilibrium of marine subsurface ecosystems [9–12].

The phylum Chloroflexi, formerly called the ‘green nonsulfur bacteria’, is a remarkably diverse, deeply branching lineage in the domain Bacteria [13]. Currently, the phylum Chloroflexi is divided phylogenetically into nine classes, including Chloroflexia [14], Anaerolineae [15], Caldilineae [15], Ktedonobacteria [16], Thermomicrobia [17], Dehalococcoidia [18], Tepidiformia [19], Thermoflexia [20] and Ardenticatenia [21]. Concomitant with the expansion of the phylum Chloroflexi by cultivation, studies utilizing cultivation-independent techniques have revealed a remarkable diversity of as-yet uncultivated microorganisms affiliated with the phylum Chloroflexi [22], indicating immeasurable novel lineages of Chloroflexi existing in nature. Despite the Chloroflexi bacteria being among the first widespread microbial lineages discovered in deep-sea environments we still lack cultured representatives (especially those with relative fast growing rate) for this group and their detailed physiological, genetic and ecological properties are currently almost completely obscure [13,23,24]. For example, until now, only basic physiological characteristics of two cultured strains of Chloroflexi with extremely slow growth rate (doubling time from 1.5 days to 19 days) from the deep-sea sediments are available [10,23], and their central metabolisms and contributions to biogeochemical processes including sulfur cycling are largely unknown.

The cycling of sulfur is one of Earth’s major biogeochemical processes and is closely related to the energy metabolism of microorganisms living in the cold seep and hydrothermal vents [25–27]. Importantly, the coupling of sulfate/sulfite reduction to oxidation of H_2_, small chain fatty acids, or other carbon compounds limits the availability of these substrates to other organisms like methanogens and alters the energetics via syntrophic interactions, and thereby impacting the methane production [25]. Given the importance of sulfur cycling in the deep biospheres, it is vital that we understand which organisms can carry out the reactions and the pathways involved [27]. Based on metagenomic sequencing results, some SAR202 members of the phylum of Chloroflexi are predicted to be sulfite-oxidizers, making them as potential key players in the sulfur cycle at the deep marine environment [28]; based on single-cell genomic sequencing results, some members of Dehalococcoidia class within the phylum Chloroflexi are demonstrated to possess diverse genes encoding dissimilatory sulfite reductase [4], suggesting that Dehalococcoidia bacteria could drive sulfite reduction and respire oxidized sulfur compounds. Together, some of the members of Chloroflexi are believed to play a previously unrecognized role in sulfur cycling, which needs to be verified with cultured representatives of Chloroflexi isolated from deep-sea environments.

In this study, we checked the abundance of Chloroflexi existing in both deep-sea cold seep and hydrothermal vents. Using an enrichment medium continuously supplemented with rifampicin pressure, we have successfully isolated a novel member of Chloroflexi, strain ZRK33, from the deep marine subsurface sediments collected from a typical cold seep in the South China Sea (1,146 m water depth). Strain ZRK33 is further to shown to be a representative of a novel class of the phylum Chloroflexi, designated as Sulfochloroflexia given that strain ZRK33 is demonstrated to assimilate sulfate and thiosulfate. Lastly, the broad distribution of diverse genes encoding key enzymes driving both sulfur assimilatory and dissimilatory reduction in the metagenome-assembled genomes from deep-sea sediments is detailed analyzed.

## Materials and methods

### Sampling and operational taxonomic units (OTUs) analysis

The deep-sea sediment samples were collected by *RV KEXUE* from a typical cold seep in the South China Sea (E 119°17’07.322’’, N 22°06’58.598’’) at a depth of approximately 1,146 m and two hydrothermal vent fields in the Okinawa Trough (E 126°53’50.247’’, N 27°47’11.096’’; E 124°22’24.86’’, N 25°15’47.438’’) in July of 2018 as described previously [26]. In order to understand the abundance of Chloroflexi phylum in the deep-sea sediments, we selected eight sedimentary samples (six cold seep samples including RPC, ZC1, ZC2, ZC3, ZC4 and ZC5 at depth intervals of 0-10, 30-50, 90-110, 150-170, 210-230 and 230-250 cm, respectively; two hydrothermal vents samples including H1 and H2 at depth intervals of 0-20 cm) for OTUs sequencing performed by Novogene (Tianjin, China). Briefly, total DNAs from these samples were extracted by the CTAB/SDS method [29] and diluted to 1 ng/µL with sterile water and used for PCR template. 16S rRNA genes of distinct regions (16S V3/V4) were amplified using specific primers (341F: 5’-CCTAYGGGRBGCASCAG and 806R: 5’-GGACTACNNGGGTATCTAAT). The PCR products were purified with a Qiagen Gel Extraction Kit (Qiagen, Germany) for libraries construction. Sequencing libraries were generated using TruSeq® DNA PCR-Free Sample Preparation Kit (Illumina, USA) following the manufacturer’s instructions. The library quality was assessed on the Qubit@ 2.0 Fluorometer (Thermo Scientific, USA) and Agilent Bioanalyzer 2100 system. The library was sequenced on an Illumina NovaSeq platform and 250 bp paired-end reads were generated. Paired-end reads were merged using FLASH (V1.2.7, http://ccb.jhu.edu/software/FLASH/) [30], which was designed to merge paired-end reads when at least some of the reads overlap with those generated from the opposite end of the same DNA fragments, and the splicing sequences were called raw tags. Quality filtering on the raw tags was performed under specific filtering conditions to obtain the high-quality clean tags [31] according to the QIIME (V1.9.1, http://qiime.org/scripts/split_libraries_fastq.html) quality controlled process. The tags were compared with the reference database (Silva database, https://www.arb-silva.de/) using UCHIME algorithm (UCHIME Algorithm, http://www.drive5.com/usearch/manual/uchime_algo.html) [32] to detect chimera sequences, and then the chimera sequences were removed [33]. Sequence analyses were performed by Uparse software (Uparse v7.0.1001, http://drive5.com/uparse/) [34]. Sequences with ≥97% similarity were assigned to the same OTUs. The representative sequence for each OTU was screened for further annotation. For each representative sequence, the Silva Database (http://www.arb-silva.de/) [35] was used based on Mothur algorithm to annotate taxonomic information.

### Metagenomic sequencing, assembly, binning and annotation

To understand the common characteristics of Chloroflexi in deep-sea environments, four cold seep sediment samples (zhu, C1, C2 and C4) and two hydrothermal vents sediment samples (H1 and H2) were selected for metagenomic analysis in BGI (BGI, China). Briefly, total DNAs from these samples (20 g each) were extracted using the Qiagen DNeasy® PowerSoil® Pro Kit (Qiagen, Hilden, Germany) and the integrity of DNA was evaluated by gel electrophoresis. Then 0.5 μg DNA of each sample was used library construction. The library was prepared with an amplification step for each sample. And then DNAs were cleaved into 50∼800 bp fragments by the Covaris E220 ultrasonicator (Covaris, Brighton, UK) and some fragments between 150∼250 bp were selected using AMPure XP beads (Agencourt, USA) and repaired using T4 DNA polymerase (Enzymatics, USA). All next-generation sequencing (NGS) was performed on the BGISEQ-500 platform (BGI, Qingdao, China) and generated 100 bp paired-end raw reads. Quality control was performed by SOAPnuke (v1.5.6) (setting: -l 20 -q 0.2 -n 0.05 -Q 2 -d -c 0 -5 0 -7 1) [36] and the clean data were assembled using MEGAHIT (v1.1.3) (setting:--min-count 2 --k-min 33 --k-max 83 --k-step 10) [37]. Assemblies of these samples were automatically binned using Maxbin2 [38], metaBAT2 [39] and Concoct [40]. MetaWRAP [41] was used to purify and organize data to generate the final bins. Finally, the completeness and contamination of metagenome-assembled genomes (MAGs) were assessed by the checkM (v1.0.18) [42]. These obtained MAGs were subsequently annotated by searching these predicted genes against KEGG (Release 87.0), NR (20180814), Swissprot (release-2017_07) and COG (update-2018_08) databases. Additionally, we utilized a custom hmmer as well as the Pfam and TIGRFAM databases to search for genes associated with sulfur metabolism using hmmsearch (e-value cut-off of 1e-20) [43].

### Enrichment and cultivation of deep-sea Chloroflexi bacteria

To enrich the Chloroflexi bacteria, deep-sea sediment samples were cultured at 28 °C for one month in an anaerobic enrichment medium (containing 1.0 g/L NH_4_Cl, 1.0 g/L NaHCO_3_, 1.0 g/L CH_3_COONa, 0.5 g/L KH_2_PO_4_, 0.2 g/L MgSO_4_.7H_2_O, 1.0 g/L yeast extract, 1.0 g/L peptone, 0.7 g/L cysteine hydrochloride, 500 µL/L 0.1 % (w/v) resazurin, pH 7.0) with 50 µg/mL rifampicin. This medium was prepared anaerobically as previously described and named ORG in this study [44]. A 50 µL enrichment culture was spread on the Hungate tubes containing ORG broth supplemented with 15 g/L agar after 10,000 times dilution. These Hungate tubes were anaerobically incubated at 28 °C for 7 days. Individual colonies were respectively picked using sterilized bamboo sticks and then cultured in the ORG broth. Strain ZRK33 was isolated and purified by repeated use the Hungate roll-tube methods for several rounds until it was considered to be axenic. The purity of strain ZRK33 was confirmed by transmission electron microscopy (TEM) and repeated partial sequencing of the 16S rRNA gene. Strain ZRK33 was preserved at -80 °C in ORG broth supplemented with 20% (v/v) glycerol.

### TEM observation

To observe the morphological characteristics of strain ZRK33, the cell suspension of fresh culture was collected at 5,000 ×*g* for 10 min and washed with Milli-Q water, and then taken by immersing copper grids coated with a carbon film for 10 min. Thereafter, the copper grids were washed for 10 min in Milli-Q water and dried for 20 min at room temperature [45]. Ultrathin-section electron microscopic observation was performed as described previously [46–48]. The sample was firstly preserved in 2.5% (v/v) glutaraldehyde for 8 h at 4 °C, washed three times with phosphate buffer saline (PBS) and then dehydrated in ethanol solutions of 30%, 50%, 70%, 90% and 100% for 10 min each time. Finally, the sample was embedded in a plastic resin. Ultrathin sections (50∼70 nm) of cells were prepared with an ultramicrotome (Leica EM UC7, Gemany), stained with uranyl acetate and lead citrate. All of these samples were examined using TEM (HT7700, Hitachi, Japan) with a JEOL JEM 12000 EX (equipped with a field emission gun) at 100 kV.

### Genome sequencing and genomic analysis

Genomic DNAs of strain ZRK33 were extracted from 2 L cells that cultured for 7 days at 28 °C. The DNA library was prepared using the Ligation Sequencing Kit (SQK-LSK109, UK), and sequenced using a FLO-MIN106 vR9.4 flow-cell for 48 h on MinKNOWN software v1.4.2 (Oxford Nanopore Technologies, UK). Whole-genome sequence determinations of strain ZRK33 were carried out with the Oxford Nanopore MinION (Oxford, UK) and Illumina MiSeq sequencing platform (San Diego, CA). A hybrid approach was utilized for genome assembly using reads from both platforms. Base-calling was performed using Albacore software v2.1.10 (Oxford Nanopore Technologies, UK). Nanopore reads were processed using protocols toolkit for quality control and downstream analysis [49]. Filtered reads were assembled using Canu version 1.8 [50] with the default parameters for Nanopore data. Finally, the genome was assembled into a single contig and was manually circularized by deleting an overlapping end.

The genome relatedness values were calculated by multiple approaches: Average Nucleotide Identity (ANI) based on the MUMMER ultra-rapid aligning tool (ANIm), ANI based on the BLASTN algorithm (ANIb), the tetranucleotide signatures (Tetra), and *in silico* DNA-DNA similarity. ANIm, ANIb and Tetra frequencies were calculated using JSpecies WS (http://jspecies.ribohost.com/jspeciesws/) [51]. The recommended species criterion cut-offs were used: 95% for the ANIb and ANIm and 0.99 for the Tetra signature. The *in silico* DNA-DNA similarity values were calculated by the Genome-to-Genome Distance Calculator (GGDC) (http://ggdc.dsmz.de/) [52]. The *is*DDH results were based on the recommended formula 2, which is independent of genome size.

### Phylogenetic analysis

The full-length 16S rRNA gene sequence (1,489 bp) of strain ZRK33 was extracted from the genome, which had been deposited in the GenBank database (accession number MN817941), and other related taxa used for phylogenetic analysis were obtained from NCBI (www.ncbi.nlm.nih.gov/). The genome tree was constructed from a concatenated alignment of 37 protein-coding genes [53] that extracted from each genome by Phylosift (v1.0.1) [54], all of which were in a single copy and universally distributed in both archaea and bacteria (Supplementary Table S1). The genomes used to construct the genome tree included both draft and finished genomes from the NCBI databases. The RpoB and EF-tu tree was constructed by using RpoB or EF-tu protein sequences, which were identified from 49 genomes using the hidden markov models (HMMs) TIGR02029 and TIGR00485 from TIGRfams (http://www.jcvi.org/cgi-bin/tigrfams/index.cgi), respectively. Phylogenetic trees were constructed by using W-IQ-TREE web server (http://iqtree.cibiv.univie.ac.at) [55] with LG+F+I+G4 model. The online tool Interactive Tree of Life (iTOL v5) [56,57] was used for editing trees.

### Growth assays of strain ZRK33

Growth assays were performed at atmospheric pressure. Briefly, 15 mL fresh strain ZRK33 culture was inoculated in 2 L Hungate bottles containing 1.5 L ORG broth supplemented with 20 mM Na_2_SO_4_, 200 mM Na_2_SO_4_, 20 mM Na_2_S_2_O_3_, 200 mM Na_2_S_2_O_3_, 1 mM Na_2_SO_3_ and 1 mM Na_2_S, respectively. Each condition had three replicates. These Hungate bottles were then anaerobically incubated at 28 °C for 12 d. Bacterial growth status was monitored by measuring the OD_600_ value every 12 h until cell growth reached the stationary phase. For the morphological observation of strain ZRK33, we took 20 µL culture that cultivated for 12 d, which was then checked and recorded under an inverted microscope (NIKON TS100, Tokyo, Japan) equipped with a digital camera. For the determination of the dynamics of the concentrations of Na_2_SO_4_ and Na_2_S_2_O_3_ in the culture, we selected three cultivation time points at 5 d, 8 d and 12 d, respectively, and each condition had three replicates. The supernatant was collected at 12,000 *g* for 10 min and diluted 80 times, and the concentrations of SO_4_^2-^ and S_2_O_3_ ^2-^ in the diluted supernatant were respectively measured by the ion chromatograph (ECO IC, Herisau, Switzerland) with an chromatographic column (Metrosep A Supp5). The column was eluted with mobile phase A (3.2 mmol/L Na_2_CO_3_) and mobile phase B (1.0 mmol/L NaHCO_3_) at 25 °C.

### Proteomic analysis

Proteomic analysis was performed by PTMBiolabs (Hangzhou, China). Briefly, strain ZRK33 was respectively cultivated in the ORG broth (set as the control group and indicated as “C”), ORG broth supplemented with 200 mM Na_2_SO_4_ (set as the experimental group and indicated as “S”) and 200 mM Na_2_S_2_O_3_ (set as the experimental group and indicated as “T”) for 8 d at 28 °C. Then the cells were collected and sonicated three times on ice using a high intensity ultrasonic processor in lysis buffer (8 M urea, 1% Protease Inhibitor Cocktail). The remaining debris was removed by centrifugation at 12,000 *g* at 4 °C for 10 min. Finally, the supernatant was collected and the protein concentration was determined with a BCA kit (Solarbio, China) according to the instructions. The detailed protocols of proteomics sequencing technology were described in the Supplementary information. The heat map was made by HemI 1.0 based on the KEGG enrichment results.

## Data availability

The raw amplicon sequencing data have been deposited to NCBI Short Read Archive (accession numbers: PRJNA675395 and PRJNA688815). The BioProject accession number of metagenome-assembled genomes (MAGs) of Chloroflexi bacteria used in this study is PRJNA667788. The full-length 16S rRNA gene sequence of *S. methaneseepsis* ZRK33 has been deposited at GenBank under the accession number MN817941. The complete genome sequence of *S. methaneseepsis* ZRK33 has been deposited at GenBank under the accession number CP051151. The mass spectrometry proteomics data have been deposited to the Proteome Xchange Consortium with the dataset identifier PXD023380.

## Results

### Chloroflexi bacteria possess high abundance in the deep-sea environments

To gain preliminary insights of Chloroflexi bacteria existing in the deep-sea environments, OTUs sequencing was firstly performed to detect the abundance of the phylum Chloroflexi present in the cold seep sediments at depth intervals of 0-10 cm (sample RPC), 30-50 cm (sample ZC1), 90-110 cm (sample ZC2), 150-170 cm (sample ZC3), 210-230 cm (sample ZC4), 230-250 cm (sample ZC5) and hydrothermal vents sediments at depth intervals of 0-20 cm from the surface to the deep layer in two different sampling sites (samples H1 and H2). As previously reported [2,6,7], the Chloroflexi group was both the second most abundant phylum in cold seep and hydrothermal vents sediments, suggesting Chloroflexi was dominant in these deep-sea regions (Figs. 1A and 1B). The proportion of Chloroflexi respectively accounted for 6.15%, 10.92%, 5.04%, 8.67% and 13.67% of the whole bacterial domain at the phylum level in samples RPC, ZC1, ZC2, H1 and H2 (Figs. 1A and 1B). Moreover, the ratio of the phylum Chloroflexi to the whole bacteria domain is much higher in the top layer than that in the bottom one, providing a preliminary hint of distribution of Chloroflexi bacteria in the deep-sea sediments. To obtain further insights into the deep-sea Chloroflexi bacteria, we analyzed the abundance of Chloroflexi members at the class level and found that Dehalococcoidia and Anaerolineae were the top two classes in the cold seep (Fig. 1C) and hydrothermal vents sediments (Fig. 1D). In particular, Dehalococcoidia class bacteria are an absolute dominant population in 7 of 8 samples, strongly suggesting the importance of this lineage in deep-sea environments.

**Fig. 1.**
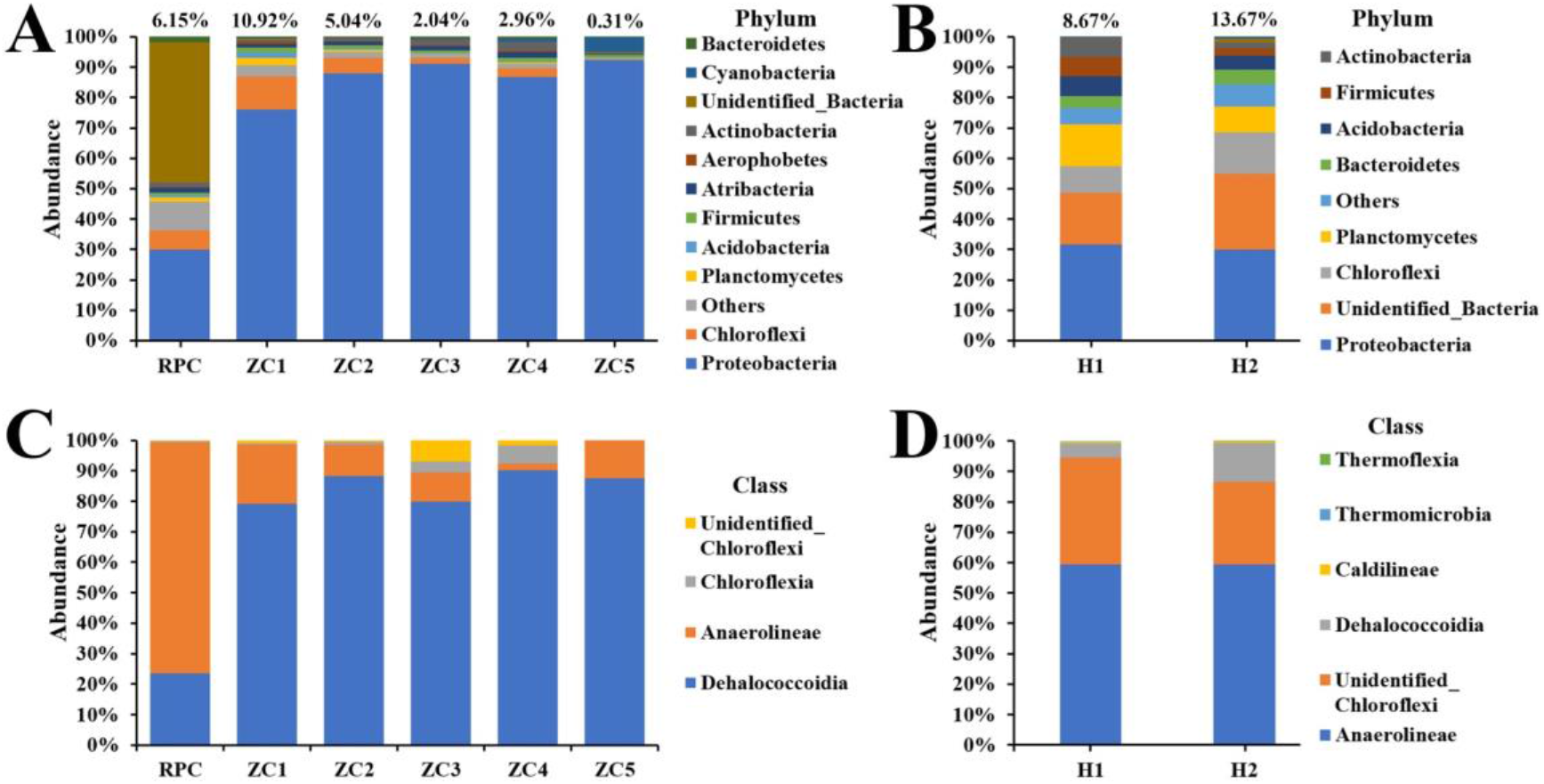
Detection of the abundance of the phylum Chloroflexi derived from the deep-sea cold seep and hydrothermal vents sediments. The community structure of six sampling sites in the cold seep sediments and two sampling sites in the hydrothermal vents sediments as revealed by 16S rRNA gene amplicon profiling. The relative abundances of operational taxonomic units (OTUs) representing different bacteria are shown at the phylum level (A and B) and class level (C and D). Panels A and C represent samples from the cold seep; Panels B and D represent samples from the hydrothermal vents.

### Cultivation and morphology of a novel Chloroflexi bacterium isolated from the deep-sea cold seep

To culture novel isolates belonging to the phylum Chloroflexi from deep-sea environments, we improved the enrichment method by using a specific medium that constantly supplemented with 50 µg/mL rifampicin, given that many members of Chloroflexi were reported to tolerate rifampicin [13,23] while most of other bacteria are sensitive to this antibiotics. Using this strategy, we anaerobically enriched the deep-sea sediment samples at 28 °C for one month. Thereafter, the enriched samples were plated on solid medium in Hungate tubes, and individual colonies with distinct morphology were picked and cultured (Fig. 2A). Excitingly, some of the cultured colonies were identified as Chloroflexi bacteria based on their 16S rRNA sequences. Among them, strain ZRK33 possessed a fast growth rate and was chosen for further study. Under TEM observation, the cells of strain ZRK33 were filamentous, generally more than 20 µm long and 0.5 -0.6 µm wide, and had no flagellum (Figs. 2B and 2C). Ultrathin sections of whole cells of strain ZRK33 revealed a cytoplasmic membrane surrounded by a cell wall surface layer (Figs. 2D and 2E). The strain did not possess a clearly visible sheath-like structure (Fig. 2D) as shown in the *Pelolinea submarina* strain MO-CFX1^T^, a typical Chloroflexi bacterium belonging to the class Anaerolineae [23]. Based on the 16S rRNA sequence of strain ZRK33, a sequence similarity calculation using the NCBI server indicated that the closest relatives of strain ZRK33 were *Anaerolinea thermophila* UNI-1 (82.75%) (class Anaerolineae), *Ornatilinea apprima* P3M-1 (82.32%) (class Anaerolineae), *Thermomarinilinea lacunifontana* SW7 (82.42%) (class Anaerolineae) and *Caldilinea aerophila* DSM 14535 (81.87%) (class Cadilineae). Recently, taxonomic thresholds based on 16S rRNA gene sequence identity values were suggested [58]: for classes, the proposed thresholds for median and minimum sequence identity values were 86.35% and 80.38%, respectively. Based on these criteria, we propose that strain ZRK33 might be a representative of a novel class-level Chloroflexi.

**Fig. 2.**
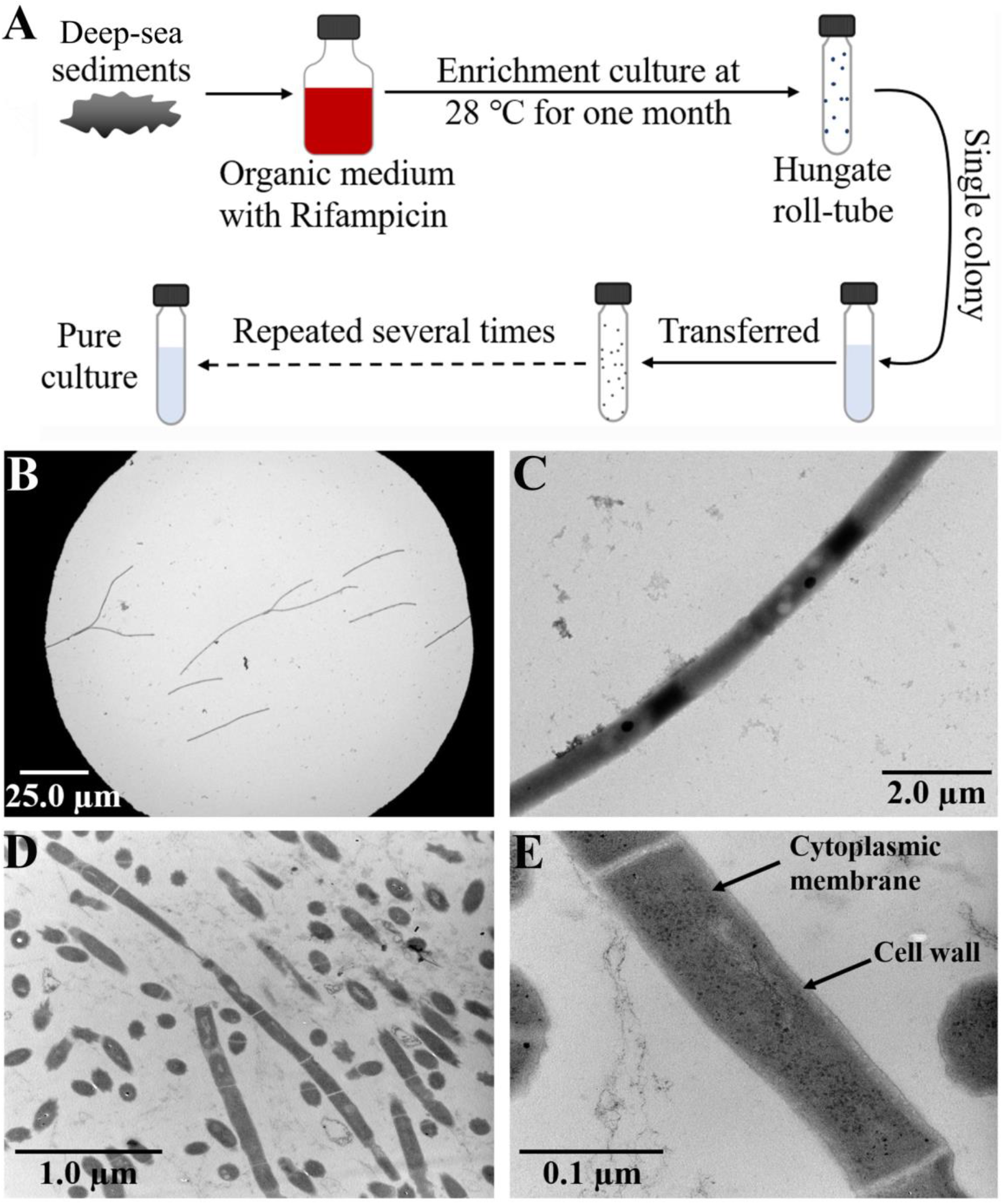
Rifampicin resistance-driven enrichment and isolation strategy of Chloroflexi bacteria. (A) Diagrammatic scheme of enrichment and isolation of Chloroflexi bacteria. (B, C) TEM observation of strain ZRK33. (D, E) TEM observation of the ultrathin sections of strain ZRK33.

### Genomic characteristics and phylogenetic analysis of strain ZRK33

To understand more characteristics of strain ZRK33, its whole genome was sequenced and analyzed. The genome size of strain ZRK33 was 5,631,885 bp with a DNA G+C content of 52.76% (Fig. 3A and Supplementary Table S2). Annotation of the genome of strain ZRK33 revealed it consisted of 4,885 predicted genes that included 55 RNA genes (6 rRNA genes, 46 tRNA genes and 3 other ncRNAs). When exploring the detailed genomic composition of strain ZRK33, we found that various genes encoding key enzymes responsible for sulfur metabolism existing in the genome of ZRK33 (Fig. 3B). And these enzymes are thought to involve in assimilatory sulfate reduction, strongly indicating that strain ZRK33 might be a representative of a novel clade that driving deep-sea sulfur cycling.

**Fig. 3.**
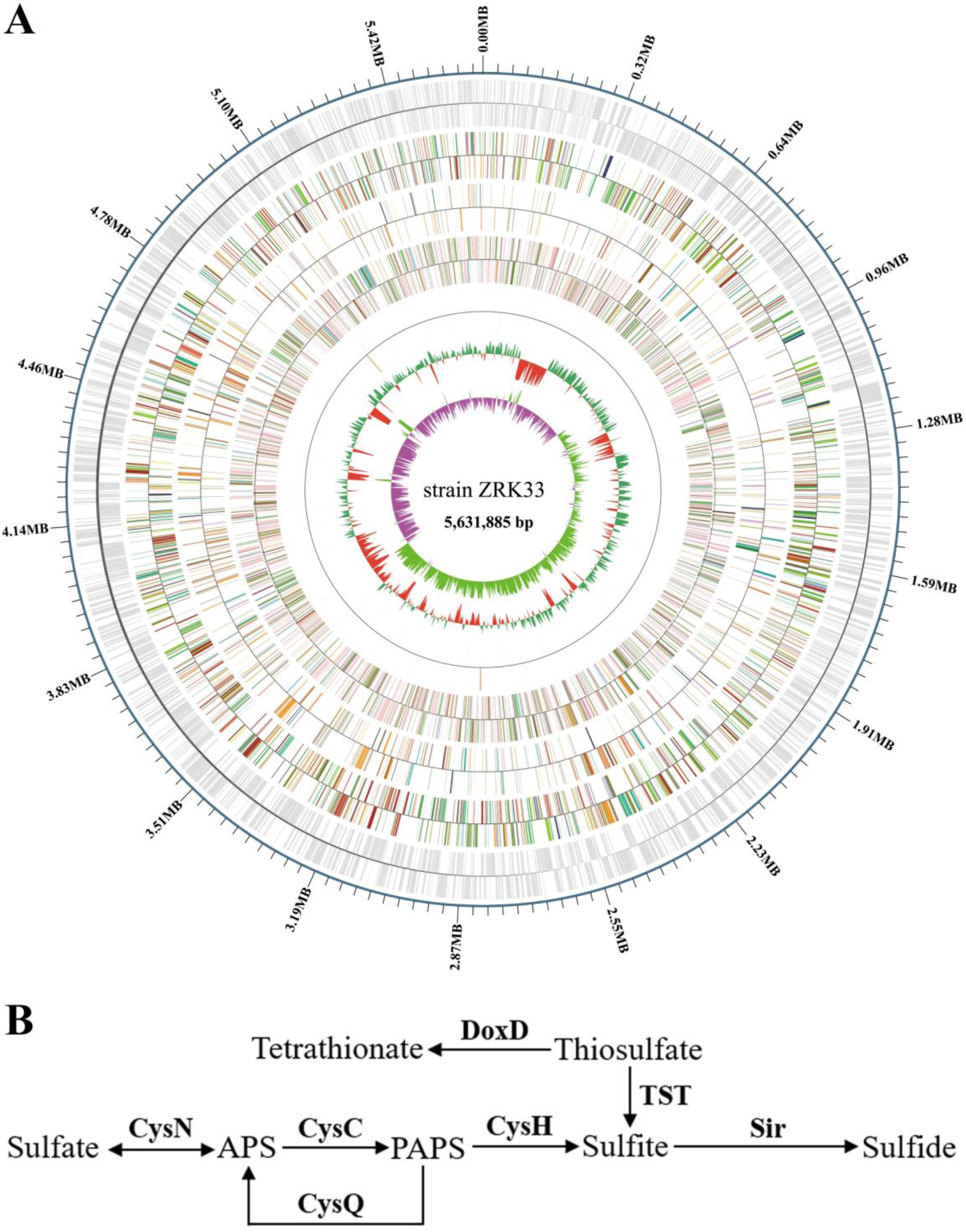
Genomic analysis of strain ZRK33. (A) Circular diagram of the genome of strain ZRK33. Rings indicate, from outside to the center: a genome-wide marker with a scale of 320 kb; forward strand genes, colored by COG category; reverse strand genes, colored by COG category; gene function annotation (COG, KEGG, GO, NR, CAZy, TCDB); RNA genes (tRNAs blue, rRNAs purple); GC content; GC skew. (B) Proposed assimilatory sulfate reduction pathway identified in the genome of strain ZRK33. Abbreviations: CysN, sulfate adenylyltransferase; Sat, sulfate adenylyltransferase; CysC, adenylyl-sulfate kinase; CysQ, 3’, 5’-bisphosphate nucleotidase; CysH, phosphoadenosine phosphosulfate reductase; Sir, sulfite reductase; TST, thiosulfate/3-mercaptopyruvate sulfurtransferase; DoxD, thiosulfate dehydrogenase (quinone) large subunit; APS, adenosine 5’-phosphosulfate; PAPS, 3’-phosphoadenosine-5’-phosphosulfate (3’-phosphoadenylylsulfate).

To further clarify the phylogenetic position of strain ZRK33, the genome relatedness values were calculated by the average nucleotide identity (ANI), *in silico* DNA-DNA similarity (*is*DDH) and the tetranucleotide signatures (Tetra), against six genomes (strain ZRK33, and five strains MO-CFX2, MO-CFX1, UNI-1, IMO-1 and P3M-1 belonging to class Anaerolineae) (Supplementary Table S3). The average nucleotide identities (ANIb) of ZRK33 with strains MO-CFX2, MO-CFX1, UNI-1, IMO-1 and P3M-1 were 64.81%, 63.06%, 63.42%, 63.41% and 63.29%, respectively. The average nucleotide identities (ANIm) of ZRK33 with MO-CFX2, MO-CFX1, UNI-1, IMO-1 and P3M-1 were 85.21%, 82.63%, 83.42%, 83.15% and 83.23%, respectively. The tetra values of ZRK33 with MO-CFX2, MO-CFX1, UNI-1, IMO-1 and P3M-1 were 0.48145, 0.67572, 0.64677, 0.65234 and 0.65126. Based on digital DNA-DNA hybridization employing the Genome-to-Genome Distance Calculator GGDC, the in silico DDH estimates for ZRK33 with MO-CFX2, MO-CFX1, UNI-1, IMO-1 and P3M-1 were 23.30%, 24.20%, 20.40%, 21.60% and 23.80%, respectively. These results together demonstrated the genome of strain ZRK33 to be obviously below established ‘cut-off’ values (ANIb: 95%, ANIm: 95%, isDDH: 70%, Tetra: 0.99) for defining bacterial species, suggesting strain ZRK33 represents a novel taxon within the phylum Chloroflexi as currently defined.

To further confirm the taxonomic status of strain ZRK33, we performed the phylogenetic analyses with 16S rRNA genes from all cultured Chloroflexi representatives, some uncultured SAR202 representatives and other uncultured Chloroflexi bacteria. The maximum likelihood tree of 16S rRNA placed strain ZRK33 as a sister of the strain MO-CFX2, which together formed a distinct cluster separating from other classes of the phylum Chloroflexi (Fig. 4). Furthermore, the genome tree also placed the novel clade as a sister of the Anaerolineae class belonging to the phylum Chloroflexi (Supplementary Figure S1). The phylogenetic analysis of strain ZRK33 using the beta subunit of RNA polymerase (RpoB), which also showed that the novel clade formed a separate branch from the Anaerolineae class (Supplementary Figure S2). More importantly, the broader phylogeny of elongation factor Tu (EF-Tu) supported the placement of the novel clade within the phylum Chloroflexi (Supplementary Figure S3). Based on phylogenetic, genomic and phenotypic characteristics, we proposed that strain ZRK33 together with strain MO-CFX2 (previously classified as a representative of a novel order of class Anaerolineae) were classified as the type strains of a new class of the phylum Chloroflexi. Given the broad distribution of genes associated with sulfur metabolism in the genome of strain ZRK33 (Fig. 3B) and its significant potential involved in sulfur cycling, we propose Sulfochloroflexia classis nov., Sulfochloroflexales ord. nov., Sulfochloroflexaceae fam. nov. and *Sulfochloroflexus methaneseepsis* gen. nov.

**Fig. 4.**
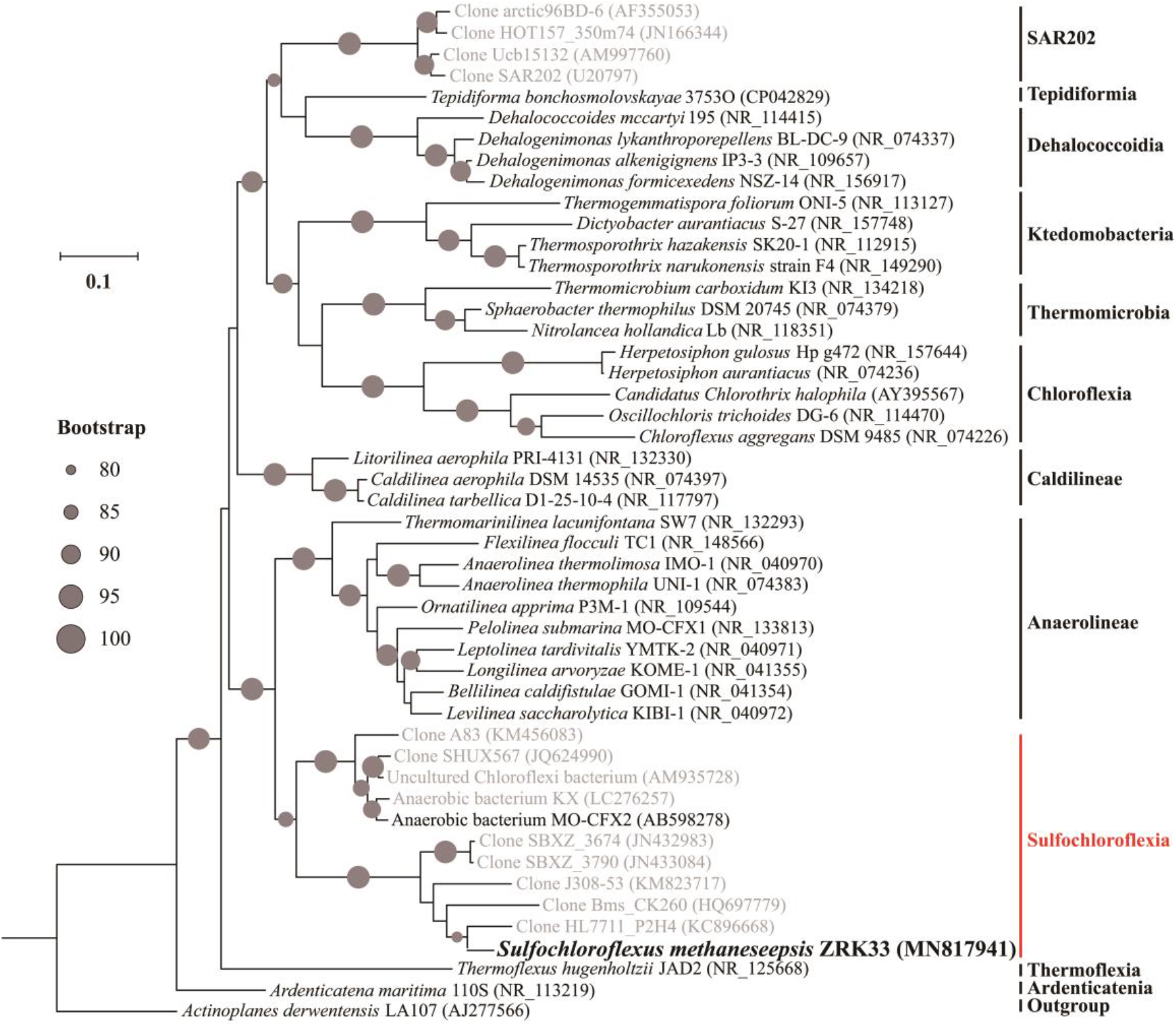
Phylogenetic analysis of *Sulfochloroflexus methaneseepsis* ZRK33. Phylogenetic placement of strain ZRK33 within the phylum Chloroflexi based on almost complete 16S rRNA gene sequences. The tree is inferred and reconstructed under the maximum likelihood criterion and bootstrap values (%) > 80 are indicated at the base of each node with the black dots (expressed as percentages of 1,000 replications). Names indicated with grey color in quotation represent taxa that are not yet validly published. All sequences are labeled with their NCBI accession numbers. The 16S rRNA gene sequence of *Actinoplanes derwentensis* LA107^T^ is used as an outgroup. Bar, 0.1 substitutions per nucleotide position.

### Description of *Sulfochloroflexus* gen. nov. and *Sulfochloroflexus methaneseepsis* sp. nov

*Sulfochloroflexus* (Sul.fo’ch.lo.ro.fle.xus. N.L. fem. pl. n. Sulfo, sulfur; N.L. masc. *chloroflexus* a bacterial genus; N.L. masc. n. *Sulfochloroflexus, chloroflexus* sulfur loving. Facultatively anaerobic, mesophilic, neutrophilic and moderately halophilic (Supplementary Table S2). Cells are non-motile. Gram-staining reaction is negative. The phylogenetic position is in the family Sulfochloroflexaceae, order Sulfochloroflexales, class Sulfochloroflexia of the phylum Chloroflexi. The type species is *Sulfochloroflexus methaneseepsis*.

*Sulfochloroflexus methaneseepsis* (me.th.ane’seep.sis. L. gen. pl. n. *methaneseepsis* of the deep-sea methane seeps). Cells are generally more than 20 µm long and 0.5-0.6 µm wide, filamentous, facultatively anaerobic and have no flagellum. From the sole carbon source utilization test, growth is stimulated by arabinose, fructose, glucose, galactose, mannose, ribose, fumarate, pyruvate and peptone. Growing at pH values of 6.0-8.0 (optimum, pH 7.0). The temperature range for growth is 28-32 °C with an optimum at 28 °C. Growth occurs at NaCl concentrations between 0.0-5.0% with optimum growth at 3.0% NaCl. Containing significant proportions (>10 %) of the cellular fatty acids C_16:0_, C_15:0_2-OH, C_17:1_*ω*6c and C_18:1_*ω*7c. The type strain, ZRK33^T^, was isolated from the sediment of deep-sea cold seep, P.R. China. The DNA G+C content of the type strain is 52.76%.

The detailed descriptions of other levels of Sulfochloroflexia were shown in the Supplementary information.

### *S. methaneseepsis* ZRK33 assimilates sulfate and thiosulfate for growth

Given that strain ZRK33 had a complete set genes of assimilatory sulfate reduction and it was isolated from the deep-sea cold seep where is rich of different sulfur-containing compounds [25,26], thus, we tested the effects of different sulfur-containing inorganic substances (including Na_2_SO_4_, Na_2_SO_3_, Na_2_S_2_O_3_, Na_2_S) on the growth of *S. methaneseepsis* ZRK33. The results showed that the supplement of high concentration (200 mM) of Na_2_SO_4_ and Na_2_S_2_O_3_ could significantly promote the growth of strain ZRK33 (Figs. 5A and B). While low concentration of Na_2_SO_4_ and Na_2_S_2_O_3_ (20 mM) had no evident effects on the growth of strain ZRK33 (Supplementary Fig. S4), indicating this bacterium is only sensitive to high concentrations of Na_2_SO_4_ and Na_2_S_2_O_3_. Meanwhile, it is noting that the concentrations of Na_2_SO_4_ and Na_2_S_2_O_3_ were respectively decreased from 200 mM to 120 mM and 140 mM along with the growth of strain ZRK33 for 12 d, suggesting that strain ZRK33 could effectively metabolize Na_2_SO_4_ and Na_2_S_2_O_3_ (Figs. 5A and B). Moreover, the average length of filamentous cells of strain ZRK33 became apparently longer in the medium supplemented with 200 mM Na_2_SO_4_ (Fig. 5D) or Na_2_S_2_O_3_ (Fig. 5E) than that in the control group (Fig. 5C), strongly suggesting that ZRK33 could assimilate Na_2_SO_4_ and Na_2_S_2_O_3_ and thereby generating extra energy for growth. In comparison, the supplement of very low concentration (1 mM) of Na_2_SO_3_ and Na_2_S inhibited the growth of strain ZRK33 (Supplementary Fig. S4), indicating that SO_3_^2-^ and S^2-^ were harmful sulfur-containing compounds against the growth of strain ZRK33.

**Fig. 5.**
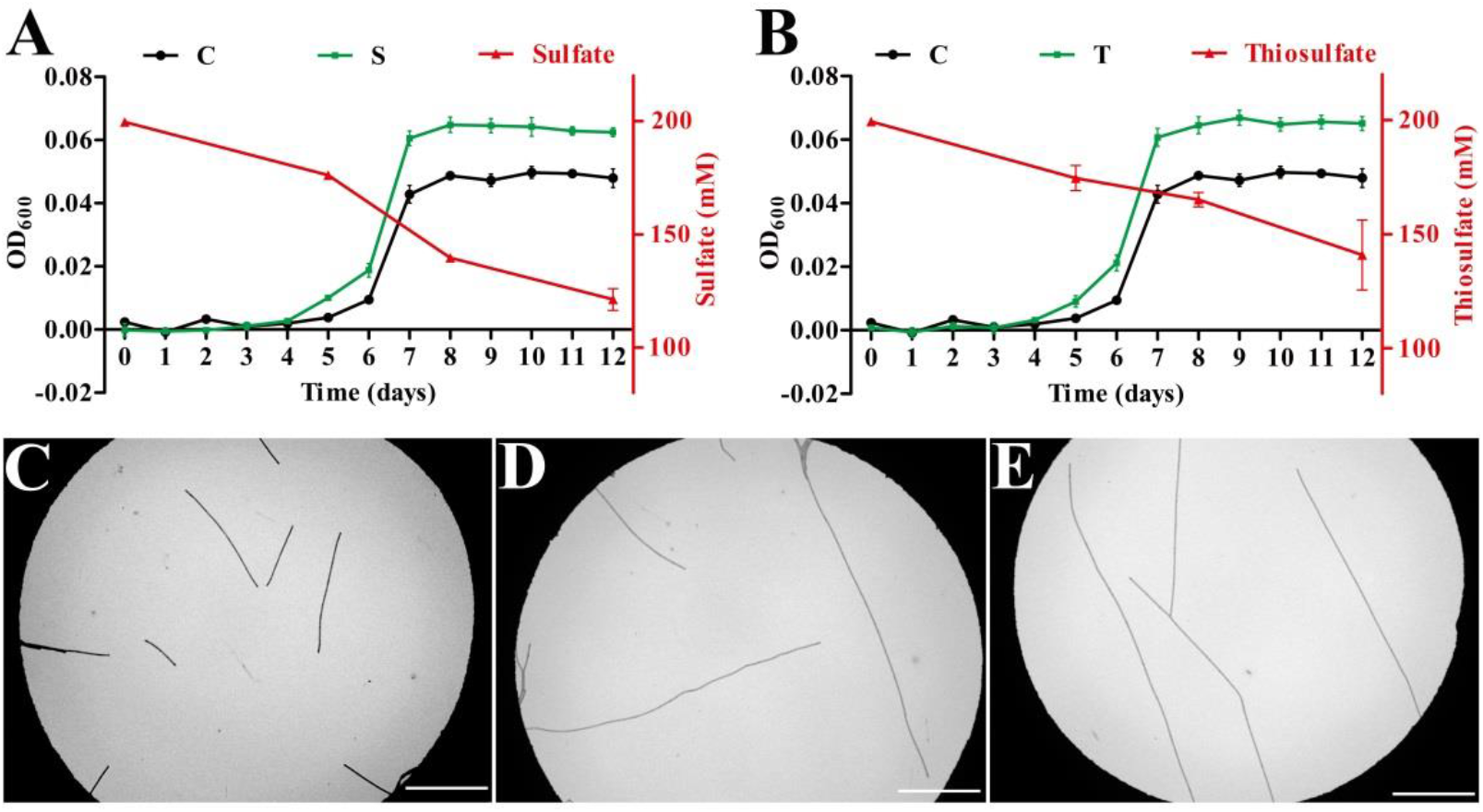
Verification of sulfate and thiosulfate assimilation in *S. methaneseepsis* ZRK33. (A) Growth assay and sulfate metabolization of strain ZRK33 cultured in the medium supplemented without or with 200 mM Na_2_SO_4_. (B) Growth assay and thiosulfate metabolization of strain ZRK33 cultured in the medium supplemented without or with 200 mM Na_2_S_2_O_3_. “C” indicates the control group, where strain ZRK33 was cultured in the medium supplemented without extra Na_2_SO_4_ or Na_2_S_2_O_3_; “S” indicates the sulfate-treated group, where strain ZRK33 was cultured in the medium supplemented with 200 mM Na_2_SO_4_; “T” indicates the thiosulfate-treated group, where strain ZRK33 was cultured in the medium supplemented with 200 mM Na_2_S_2_O_3_. The black lines represent the growth curves of the control group; the green lines represent the growth curves of experimental groups; the red lines represent the variation tendency of concentrations of Na_2_SO_4_ or Na_2_S_2_O_3_. (C) TEM observation of strain ZRK33 that cultured in the ORG medium. (D) TEM observation of strain ZRK33 that cultured in the ORG medium supplemented with 200 mM Na_2_SO_4_. (E) TEM observation of strain ZRK33 that cultured in the ORG medium supplemented with 200 mM Na_2_S_2_O_3_. The bar is 20 µm in the panels C, D and E.

### Proteomic analyses of sulfur metabolism in *S. methaneseepsis* ZRK33

To better describe the sulfur metabolism of *S. methaneseepsis* ZRK33, we performed the proteomic analysis of strain ZRK33 that cultured in the medium amended with or without Na_2_SO_4_/Na_2_S_2_O_3_ to explore the underlying mechanism of growth promotion, given that ZRK33 could effectively assimilate Na_2_SO_4_/Na_2_S_2_O_3_ for its growth. The results showed that the expression of sulfurtransferase (TST), sulfatase-like hydrolase and cysteine desulfurase-like protein were obviously up-regulation compared with the control group, which were associated with sulfur metabolism (Fig. 6A). In particular, TST is a key enzyme catalyzing S_2_O_3_^2-^ to SO_3_^2-^ and thereby joining into sulfur assimilation (Fig. 3B), and it was significantly up-regulated in the presence of high concentrations of Na_2_SO_4_/Na_2_S_2_O_3_, especially Na_2_S_2_O_3_. Surprisingly, the expressions of other proteins associated with assimilatory sulfate reduction were not significantly up-regulated in experimental groups, partly due to the single sampling time point that might miss the exact time to detect the up-regulation of key proteins associated with sulfur metabolism. Alternatively, the expressions of many proteins associated with organic matter metabolisms toward energy production were evidently up-regulated, including amino acids and sugar ABC transporters (Fig. 6B), saccharides/peptides/amino acids degradation (Fig. 6C), and energy production (Fig. 6D). Correspondingly, the expressions of almost all genes involved in EMP glycolysis were also significantly up-regulated (Supplementary Figures S5). Thus, we speculated that metabolization of sulfate and thiosulfate by strain ZRK33 may accelerate the hydrolysis and uptake of saccharides and other organic matter and thereby synthesizing energy to promote the growth [48]. Combining the results of catalyzing of sulfate and thiosulfate to other formations (Figs. 5C and 5D), we believe that strain ZRK33 possesses a capability to assimilate inorganic sulfur-containing compounds (e.g. sulfate and thiosulfate) that ubiquitously existing in the deep-sea environments and thereby contributing to the deep-sea sulfur cycling to some extent.

**Fig. 6.**
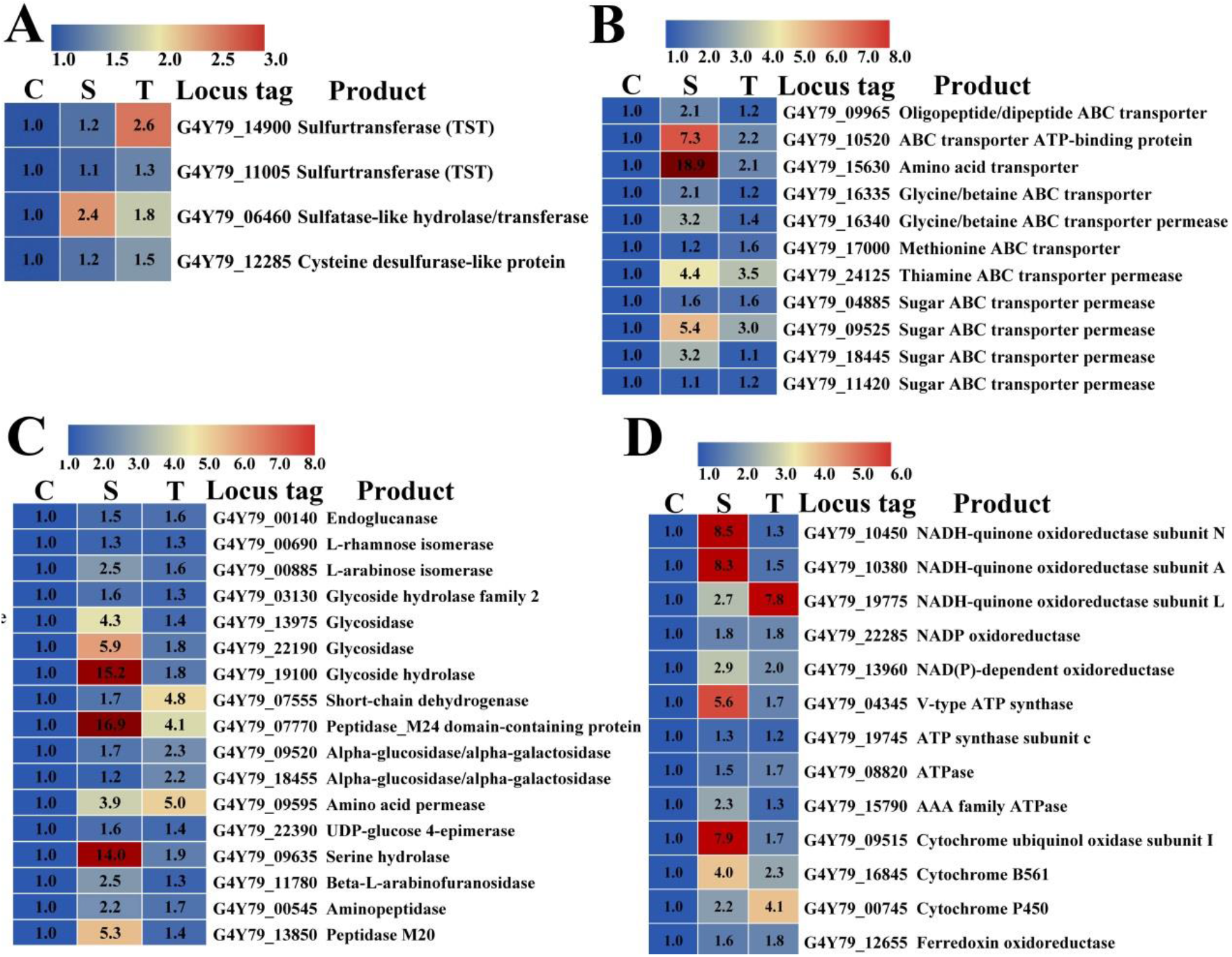
Proteomic analysis of *S. methaneseepsis* ZRK33 cultured in the medium supplemented with sulfate and thiosulfate. (A) Proteomics based heat map showing all up-regulated proteins associated with sulfur metabolism. (B) Proteomics based heat map showing all up-regulated proteins associated with amino acids and sugar transporters. (C) Proteomics based heat map showing all up-regulated proteins associated with saccharides/amino acids/peptides hydrolases. (D) Proteomics based heat map showing all up-regulated proteins associated with energy production. “C” indicates the control group, where strain ZRK33 was cultured in the medium supplemented without extra Na_2_SO_4_ or Na_2_S_2_O_3_; “S” indicates the sulfate-treated group, where strain ZRK33 was cultured in the medium supplemented with 200 mM Na_2_SO_4_; “T” indicates the thiosulfate-treated group, where strain ZRK33 was cultured in the medium supplemented 200 mM Na_2_S_2_O_3_.

Based on the combination of proteomic, genomic and physiological characteristics, we propose a model towards central metabolic traits of strain ZRK33 (Fig. 7). In this model, central metabolisms including EMP glycolysis, oxidative pentose phosphate pathway, TCA cycle (tricarboxylic acid cycle), assimilatory sulfate reduction, urea cycle and electron transport system are shown. All the above items are closely related to the energy production in strain ZRK33. Briefly, strain ZRK33 contains a number of genes related to ABC transporters of amino acids, peptides and sugar, which could transport these organic matters into the cell to participate in EMP glycolysis and oxidative pentose phosphate pathway. These processes eventually drive the formation of pyruvate and acetyl-CoA, which enter the TCA cycle to produce energy for the growth of strain ZRK33. Of note, sulfate and thiosulfate could be converted to cysteine and thereby entering the pyruvate synthesis pathway through the assimilatory sulfate reduction, which might promote the saccharides degradation and utilization via some unknown mechanisms. Moreover, strain ZRK33 could fix nitrogen and carbon dioxide to involve in the urea cycle, and corresponding metabolites can join into the TCA cycle for energy generation. Meanwhile, the F-type ATP synthase, cytochrome bd ubiquinol oxidase and H^+^-transporting NADH: Quinone oxidoreductase required for energy production are also present in the genome of strain ZRK33. Overall, ***S***. *methaneseepsis* ZRK33 is a representative of a novel clade of the phylum Chloroflexi that possessing diverse metabolic pathways for energy production, providing an evidence that Chloroflexi members are a group of high-abundance bacteria ubiquitously distributed in different environments.

**Fig. 7.**
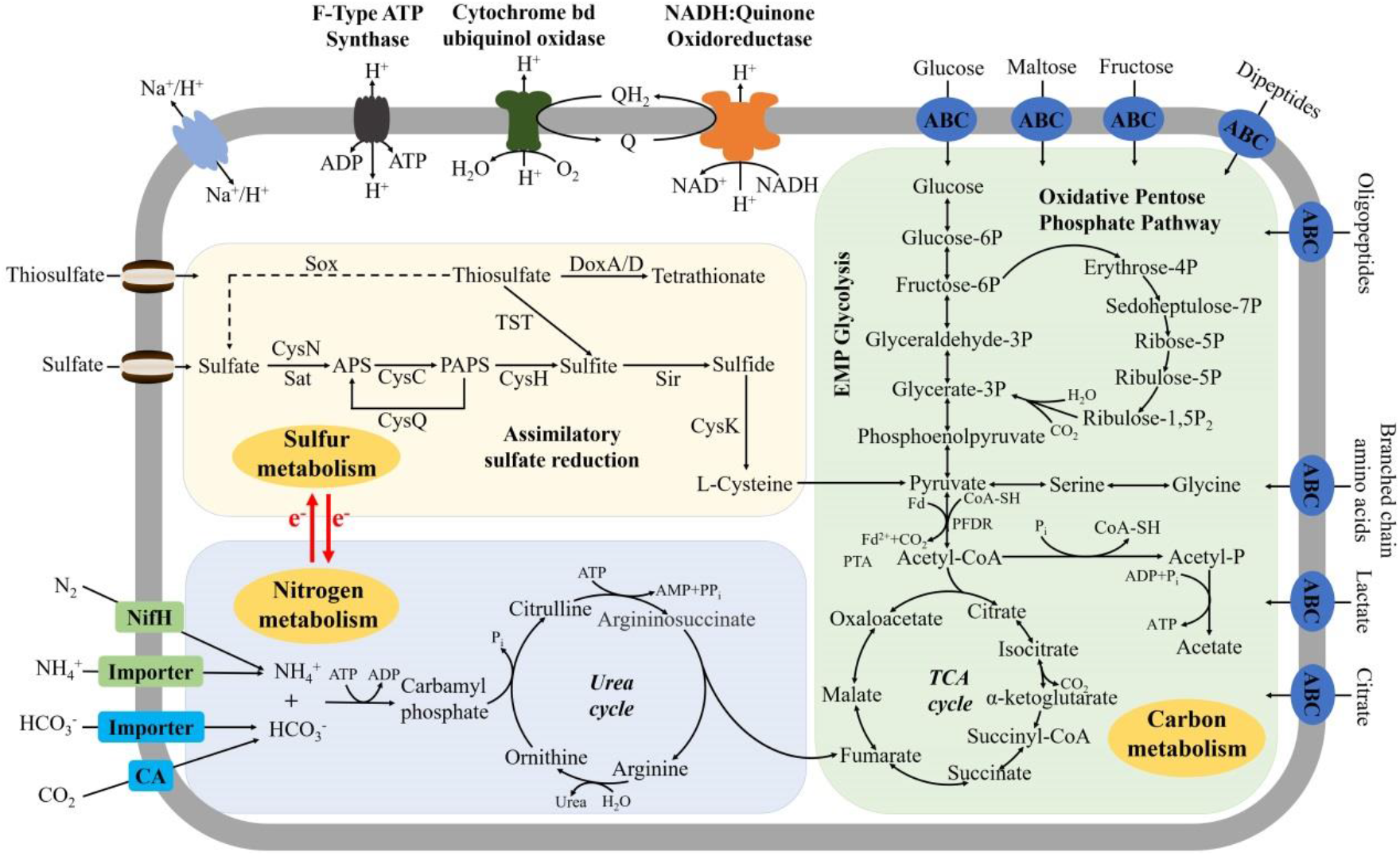
Muti-omics based central metabolisms model of *S. methaneseepsis* ZRK33. In this model, three central metabolic pathways (associated with carbon, sulfur and nitrogen cyclings) including EMP glycolysis, oxidative pentose phosphate pathway, TCA cycle, urea cycle, assimilatory sulfate reduction and some electron transport systems are shown and highlighted with different colors. All the above items are closely related to the energy production in *S. methaneseepsis* ZRK33. Abbreviations: TCA, tricarboxylic acid cycle; Urea, urea cycle; ATP, 5’-Adenylate triphosphate; ADP, adenosine diphosphate; AMP, adenosine monophosphate; CA, carbonic anhydrase; NifH, nitrogenase iron protein; Q, quinone; QH_2_, ubiquinone; CysN, sulfate adenylyltransferase; Sat, sulfate adenylyltransferase; CysC, adenylyl-sulfate kinase; CysQ, 3’, 5’-bisphosphate nucleotidase; CysH, phosphoadenosine phosphosulfate reductase; Sir, sulfite reductase; CysK, cysteine synthase; TST, thiosulfate/3-mercaptopyruvate sulfurtransferase; DoxA, thiosulfate dehydrogenase (quinone) small subunit; DoxD, thiosulfate dehydrogenase (quinone) large subunit; Sox, L-cysteine S-thiosulfotransferase.

### Wide distribution of assimilatory and dissimilatory sulfate reduction pathways in the deep-sea Chloroflexi bacteria

To evaluate the contribution of Chloroflexi bacteria to the deep-sea sulfur cycling, we further analyzed the distribution of genes encoding key enzymes responsible for both assimilatory (Fig. 8A) and dissimilatory (Fig. 8B) sulfate reduction in 27 metagenome-assembled genomes (MAGs) of Chloroflexi bacteria derived from both deep-sea cold seep and hydrothermal vents sediments. Through a thorough analysis of 27 MAGs, we found that diverse genes encoding key enzymes in charge of assimilatory and dissimilatory sulfate reduction, including adenylyl-sulfate kinase (CysC), 3’, 5’-bisphosphate nucleotidase (CysQ), sulfate adenylyltransferase (CysN), anaerobic sulfite reductase (AsrA, AsrB and AsrC) and dissimilatory sulfite reductase (DsrA and DsrB), were widely distributed in both cold seep and hydrothermal vents derived MAGs (Fig. 8C). Of note, genes encoding AsrA and AsrB were present in most MAGs, however, genes encoding DsrA and DsrB only broadly existed in the hydrothermal vents-derived MAGs (Fig. 8C). DsrA and DsrB are typical symbols of microbes mediating dissimilatory sulfate reduction [4]. Therefore, we propose dissimilatory sulfate reduction might be often adopted by the members of Chloroflexi in the hydrothermal vents. Nonetheless, Chloroflexi bacteria should be important participants in sulfur cycling in the deep-sea environments, given their high abundance in both cold seep and hydrothermal vents.

**Fig. 8.**
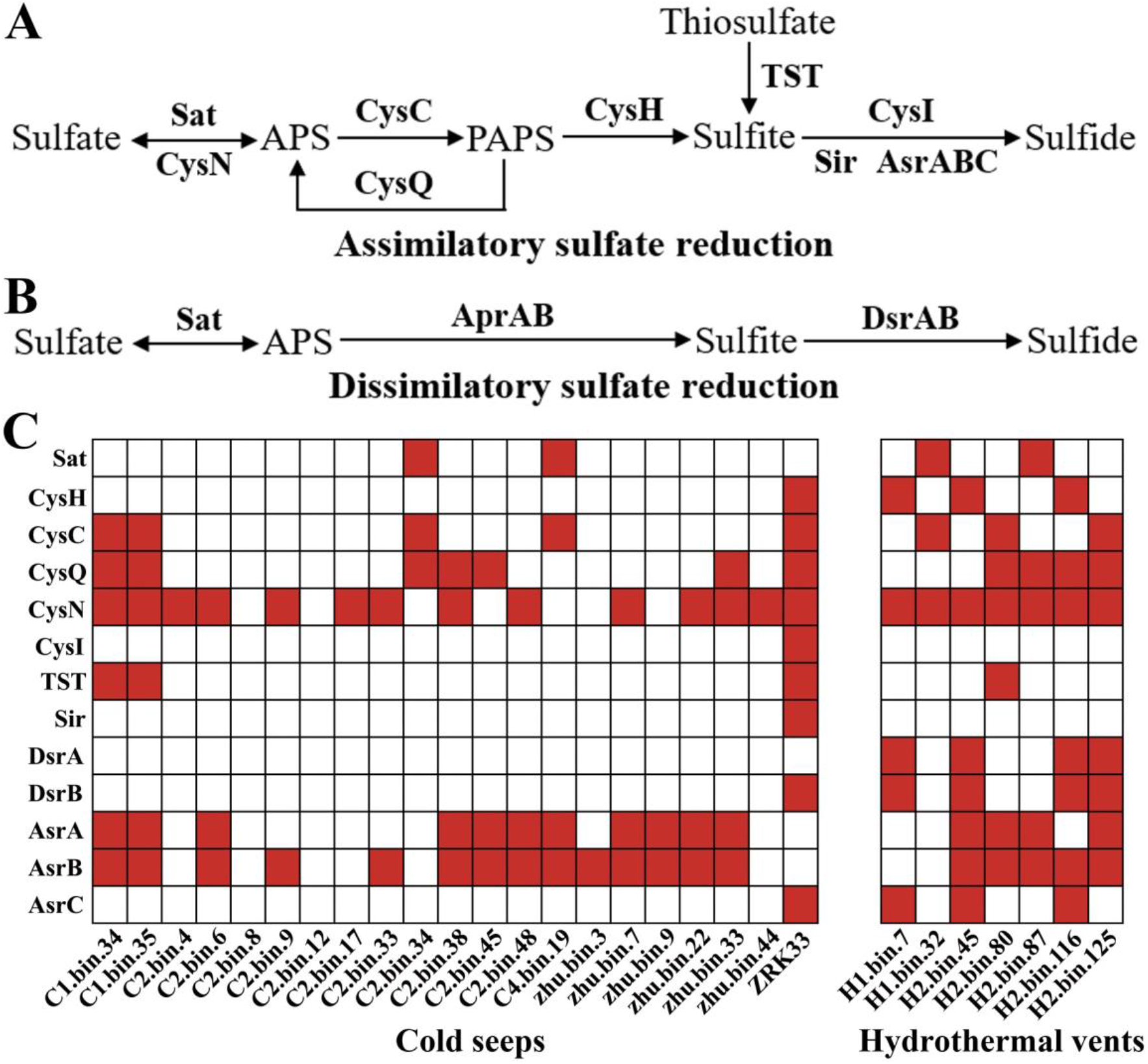
Broad distribution of genes encoding key enzymes driving assimilatory and dissimilatory sulfate reduction pathways in the metagenome-assembled genomes (MAGs) of Chloroflexi bacteria derived from deep-sea cold seep and hydrothermal vents sediments. (A) Typical pathway of assimilatory sulfate reduction existing in bacteria. (B) Typical pathway of dissimilatory sulfate reduction existing in bacteria. (C) Distribution of genes encoding key enzymes involved in assimilatory and dissimilatory sulfur metabolisms in deep-sea Chloroflexi MAGs and strain ZRK33. The presence of enzymes involved in the sulfur metabolic pathway is indicated by using red colored rectangles. Sat, sulfate adenylyltransferase; CysN, sulfate adenylyltransferase; CysC, adenylyl-sulfate kinase; CysQ, 3’, 5’-bisphosphate nucleotidase; CysH, phosphoadenosine phosphosulfate reductase; Sir, sulfite reductase; AsrA, AsrB and AsrC, anaerobic sulfite reductases; CysI, sulfite reductase (NADPH) hemoprotein beta-component; TST, thiosulfate/3-mercaptopyruvate sulfurtransferase; AprA and AprB, adenylylsulfate reductase; DsrA and DsrB, dissimilatory sulfite reductase.

## Discussion

Microorganisms in the deep marine subsurface sediments represent a large unexplored biosphere, exploring and resolving their metabolisms are essential to understand the global biogeochemical cycles [59–61]. Despite the global importance of these microorganisms, deep-sea sediments are among the least understood environments, partly due to the difficulty of sampling as well as the complexity of inhabiting communities [59]. With this, the majority of deep-sea microbial diversity remains uncultured, hampering a more thorough understanding of their unique biology [3,59,62]. One of the striking characteristics of these uncultured lineages is that most of them are dominant population, for example, the large proportion of uncultured microbes was estimated to make up more than 75% of sediment genera [63]. Therefore, it is crucial to increase our capability for bringing microorganisms from the environment into culture [60], which will advance our understanding of their global biogeochemical cycles [48]. Among the uncultured majorities, the phylum Chloroflexi is ubiquitous and often abundant in sediments, soils and wastewater treatment systems, as well as in deep-sea extreme environments [64].

Indeed, our OTUs sequencing results clearly show that the abundance of the phylum Chloroflexi is both the second most in the domain Bacteria that living in the cold seep and hydrothermal vents sediments (Figs. 1A and 1B). Although Chloroflexi bacteria are widespread on Earth, the puzzling thing is the extreme difficulty to culture Chloroflexi members from various environments, leading to a poor understanding with regard to their unique metabolisms that endowing them with tremendous vitality. Therefore, it is an urgent need to obtain more uncultivated isolates for better resolving their diversity and ecological roles, especially from the deep-sea environments given their potentials to participate in sulfur cycling [4,28]. When we looked through the previously reported characteristics of Chloroflexi isolates, one of the striking features attracted our attention: most cultured Chloroflexi could tolerate a high concentration of rifampicin (50 µg/mL) [13,23]. It is known that rifampicin is an effective inhibitor of DNA-dependent RNA polymerase and inhibits the growth of many bacteria [23]. Therefore, in the present study, we developed an effective enrichment method by keeping a constant rifampicin pressure in the enrichment and isolation medium (Fig. 2A). Indeed, we successfully obtained a novel Chloroflexi isolate, strain ZRK33, from the cold seep samples (Fig. 2). Strikingly, strain ZRK33 possessed a very fast growth rate (4 h for doubling time) compared to other reported Chloroflexi isolates (6 h to 19 days for doubling time) (Supplementary Table S2), providing a great advantage for us to promptly perform various assays. Overall, we strongly recommend the researchers to use rifampicin as a selection pressure to enrich and culture novel isolates of Chloroflexi in the future.

Additionally, we proposed strain ZRK33 as a representative of a novel class of the phylum Chloroflexi. The reasons are as following: (1) strain ZRK33 showed only ∼82% 16S rRNA gene identity with other cultured isolates, which meets the proposed thresholds for median (86.35%) and minimum (80.38%) sequence identity values to build a novel class [58]; (2) the phylogenetic analyses based on the genome (Supplementary Figure S1), beta RpoB (Supplementary Figure S2) and EF-Tu (Supplementary Figure S3) all support the classification of strain ZRK33 as the type strain of a new class; (3) strain ZRK33 is facultatively anaerobic, however, the Anaerolineae class bacteria are obligately anaerobic (Supplementary Table S2)[13], even though the novel clade shows the highest identity with the Anaerolineae class. Taken together, we propose strain ZRK33 together with *Aggregatilinea lenta* MO-CFX2^T^ to represent a novel class of Chloroflexi phylum, though strain MO-CFX2^T^ was thought to be a representative of a novel order of the class Anaerolineae [13].

Notably, we find that strain ZRK33 contains a complete set of genes associated with assimilatory sulfate reduction (Fig. 3B), providing potentials to involve into sulfur cycling. Therefore, we name this novel isolate as *Sulfochloroflexus methaneseepsis* ZRK33, which belonging to Sulfochloroflexia classis nov., Sulfochloroflexales ord. nov., Sulfochloroflexaceae fam. nov.. The cycling of sulfur is a dominant metabolism pathway for the marine subsurface microorganisms [26,65], and deep-sea Chloroflexi bacteria were predicted to respire oxidized sulfur compounds [4] and metabolize multiple organosulfur compounds [28] based on metagenomics data. However, to date, no studies based on the pure culture have verified that Chloroflexi members indeed drive sulfur cycling of deep-sea environments. Take advantage of pure cultivation of *S. methaneseepsis* ZRK33, we verified its actual involvement of both sulfate and thiosulfate assimilatory processes (Fig. 5), and the sulfur assimilatory greatly facilitates the growth and morphogenesis of ZRK33 via promoting the transport and metabolization of saccharides and other organic matter (Fig. 6). However, strain ZRK33 only responds to high concentrations of sulfate and thiosulfate (200 mM), given the high concentrations of different sulfur-containing compounds [25,26] existing in the cold seep and some microbes possessing a capability to enrich sulfur-containing compounds (such as elemental sulfur and polysulfide [26,66]), we speculate this phenomenon is possible to happen in the deep-sea cold seep sediments.

Most importantly, large portion of the genes associated with assimilatory or dissimilatory sulfate reduction are widely distributed in the Chloroflexi MAGs derived from deep-sea cold seep and hydrothermal vents (Fig. 8), which strongly suggesting Chloroflexi bacteria are key players in the sulfur cycling of deep biosphere. In combination with the reports that the other two Chloroflexi lineages (SAR202 group and Dehalococcoidia class) possessing potentials to drive sulfur metabolizations, it is reasonable to affirm the phylum Chloroflexi greatly contributes to the ocean sulfur cycling. Actually, we tried to check the metabolisms of strain ZRK33 that cultured in the deep-sea cold seep as performed previously [48], unfortunately, the cells of strain ZRK33 were invaded by some unknown microbes that leading the failure of *in situ* test toward its involvement of sulfur cycling. We are improving the experimental apparatus and procedure, which will greatly benefit us to check the central metabolisms of strain ZRK33 *in situ* in the near future.

## Acknowledgements

This work was funded by the Major Research Plan of the National Natural Science Foundation (Grant No. 92051107), China Ocean Mineral Resources R&D Association Grant (Grant No. DY135-B2-14), Key Deployment Projects of Center of Ocean Mega-Science of the Chinese Academy of Sciences (Grant No. COMS2020Q04), Strategic Priority Research Program of the Chinese Academy of Sciences (Grant No. XDA22050301), National Key R and D Program of China (Grant No. 2018YFC0310800), the Taishan Young Scholar Program of Shandong Province (tsqn20161051), and Qingdao Innovation Leadership Program (Grant No. 18-1-2-7-zhc) for Chaomin Sun. This study is also funded by the Open Research Project of National Major Science & Technology Infrastructure (*RV KEXUE*) (Grant No. NMSTI-KEXUE2017K01).

## Author contributions

RZ and CS conceived and designed the study; RZ conducted most of the experiments; RL, YS and GL collected the samples from the deep-sea cold seep; RC helped to analyze the metagenomes; RZ and CS lead the writing of the manuscript; all authors contributed to and reviewed the manuscript.

## References

1. Kallmeyer J, Pockalny R, Adhikari RR, Smith DC, D’Hondt S. Global distribution of microbial abundance and biomass in subseafloor sediment. P Natl Acad Sci USA. (2012); 109: 16213–16216.

2. Parkes RJ, Cragg BA, Wellsbury P. Recent studies on bacterial populations and processes in subseafloor sediments: A review. Hydrogeol J. (2002); 10: 346–346.

3. Inagaki F, Nunoura T, Nakagawa S, Teske A, Lever M, Lauer A, et al. Biogeographical distribution and diversity of microbes in methane hydrate-bearing deep marine sediments, on the Pacific Ocean Margin. P Natl Acad Sci USA. (2006); 103: 2815–2820.

4. Wasmund K, Cooper M, Schreiber L, Lloyd KG, Baker BJ, Petersen DG, et al. Single-cell genome and group-specific dsrAB sequencing implicate marine members of the class Dehalococcoidia (Phylum Chloroflexi) in sulfur sycling. mBio. (2016); 7.

5. Biddle JF, Fitz-Gibbon S, Schuster SC, Brenchley JE, House CH. Metagenomic signatures of the Peru Margin subseafloor biosphere show a genetically distinct environment. P Natl Acad Sci USA. (2008); 105: 10583–10588.

6. Blazejak A, Schippers A. High abundance of JS-1-and Chloroflexi-related Bacteria in deeply buried marine sediments revealed by quantitative, real-time PCR. FEMS Microbiol Ecol. (2010); 72: 198–207.

7. Parkes RJ, Cragg B, Roussel E, Webster G, Weightman A, Sass H. A review of prokaryotic populations and processes in sub-seafloor sediments, including biosphere:geosphere interactions. Mar Geol. (2014); 352: 409–425.

8. Fry JC, Parkes RJ, Cragg BA, Weightman AJ, Webster G. Prokaryotic biodiversity and activity in the deep subseafloor biosphere. FEMS Microbiol Ecol. (2008); 66: 181–196.

9. Speirs LBM, Rice DTF, Petrovski S, Seviour RJ. The phylogeny, biodiversity, and ecology of the Chloroflexi in activated sludge. Front Microbiol. (2019); 10.

10. Bovio P, Cabezas A, Etchebehere C. Preliminary analysis of Chloroflexi populations in full-scale UASB methanogenic reactors. J Appl Microbiol. (2019); 126: 667–683.

11. Schmitt S, Deines P, Behnam F, Wagner M, Taylor MW. Chloroflexi bacteria are more diverse, abundant, and similar in high than in low microbial abundance sponges. FEMS Microbiol Ecol. (2011); 78: 497–510.

12. Sorokin DY, Lucker S, Vejmelkova D, Kostrikina NA, Kleerebezem R, Rijpstra WIC, et al. Nitrification expanded: discovery, physiology and genomics of a nitrite-oxidizing bacterium from the phylum Chloroflexi. ISME J. (2012); 6: 2245–2256.

13. Nakahara N, Nobu MK, Takaki Y, Miyazaki M, Tasumi E, Sakai S, et al. Aggregatilinea lenta gen. nov., sp. nov., a slow-growing, facultatively anaerobic bacterium isolated from subseafloor sediment, and proposal of the new order Aggregatilineales ord. nov. within the class Anaerolineae of the phylum Chloroflexi. Int J Syst Evol Micr. (2019); 69: 1185–1194.

14. Gupta RS, Chander P, George S. Phylogenetic framework and molecular signatures for the class Chloroflexi and its different clades; proposal for division of the class Chloroflexi class. nov into the suborder Chloroflexineae subord. nov., consisting of the emended family Oscillochloridaceae and the family Chloroflexaceae fam. nov., and the suborder Roseiflexineae subord. nov., containing the family Roseiflexaceae fam. nov. Anton Leeuw Int J G. (2013); 103: 99–119.

15. Yamada T, Sekiguchi Y, Hanada S, Imachi H, Ohashi A, Harada H, et al. Anaerolinea thermolimosa sp nov., Levilinea saccharolytica gen. nov., sp nov and Leptolinea tardivitalis gen. nov., so. nov., novel filamentous anaerobes, and description of the new classes anaerolineae classis nov and Caldilineae classis nov in the bacterial phylum Chloroflexi. Int J Syst Evol Micr. (2006); 56: 1331–1340.

16. Yabe S, Aiba Y, Sakai Y, Hazaka M, Yokota A. Thermosporothrix hazakensis gen. nov., sp nov., isolated from compost, description of Thermosporotrichaceae fam. nov within the class Ktedonobacteria Cavaletti et al. 2007 and emended description of the class Ktedonobacteria. Int J Syst Evol Micr. (2010); 60: 1794–1801.

17. Garrity GM, Holt JG, Perry JJ. (2001) In Boone, D. R., Castenholz, R. W.and Garrity, G. M. (eds.), Bergey’s Manual® of Systematic Bacteriology: Volume One : The Archaea and the Deeply Branching and Phototrophic Bacteria. Springer New York, New York, NY, pp. 447–450.

18. Loffler FE, Yan J, Ritalahti KM, Adrian L, Edwards EA, Konstantinidis KT, et al. Dehalococcoides mccartyi gen. nov., sp nov., obligately organohalide-respiring anaerobic bacteria relevant to halogen cycling and bioremediation, belong to a novel bacterial class, Dehalococcoidia classis nov., order Dehalococcoidales ord. nov and family Dehalococcoidaceae fam. nov., within the phylum Chloroflexi. Int J Syst Evol Micr. (2013); 63: 625–635.

19. Kochetkova TV, Zayulina KS, Zhigarkov VS, Minaev NV, Chichkov BN, Novikov AA, et al. Tepidiforma bonchosmolovskayae gen. nov., sp. nov., a moderately thermophilic Chloroflexi bacterium from a Chukotka hot spring (Arctic, Russia), representing a novel class, Tepidiformia, which includes the previously uncultivated lineage OLB14. Int J Syst Evol Micr. (2020); 70: 1192–1202.

20. Dodsworth JA, Gevorkian J, Despujos F, Cole JK, Murugapiran SK, Ming H, et al. Thermoflexus hugenholtzii gen. nov., sp. nov., a thermophilic, microaerophilic, filamentous bacterium representing a novel class in the Chloroflexi, Thermoflexia classis nov., and description of Thermoflexaceae fam. nov. and Thermoflexales ord. nov.. Int J Syst Evol Micr. (2014); 64: 3331–3331.

21. Kawaichi S, Ito N, Kamikawa R, Sugawara T, Yoshida T, Sako Y. Ardenticatena maritima gen. nov., sp nov., a ferric iron- and nitrate-reducing bacterium of the phylum ‘Chloroflexi’ isolated from an iron-rich coastal hydrothermal field, and description of Ardenticatenia classis nov. Int J Syst Evol Micr. (2013); 63: 2992–3002.

22. Rappe MS, Giovannoni SJ. The uncultured microbial majority. Annu Rev Microbiol. (2003); 57: 369–394.

23. Imachi H, Sakai S, Lipp JS, Miyazaki M, Saito Y, Yamanaka Y, et al. Pelolinea submarina gen. nov., sp nov., an anaerobic, filamentous bacterium of the phylum Chloroflexi isolated from subseafloor sediment. Int J Syst Evol Micr. (2014); 64: 812–818.

24. Imachi H, Aoi K, Tasumi E, Saito Y, Yamanaka Y, Saito Y, et al. Cultivation of methanogenic community from subseafloor sediments using a continuous-flow bioreactor. ISME J. (2011); 5: 1913–1925.

25. Wasmund K, Mussmann M, Loy A. The life sulfuric: microbial ecology of sulfur cycling in marine sediments. Env Microbiol Rep. (2017); 9: 323–344.

26. Zhang J, Liu R, Xi SC, Cai RN, Zhang X, Sun CM. A novel bacterial thiosulfate oxidation pathway provides a new clue about the formation of zero-valent sulfur in deep sea. ISME J. (2020); 14: 2261–2274.

27. Fullerton H, Moyer CL. Comparative single-cell genomics of Chloroflexi from the Okinawa Trough deep-subsurface biosphere. Appl Environ Microb. (2016); 82: 3000–3008.

28. Mehrshad M, Rodriguez-Valera F, Amoozegar MA, Lopez-Garcia P, Ghai R. The enigmatic SAR202 cluster up close: shedding light on a globally distributed dark ocean lineage involved in sulfur cycling. ISME J. (2018); 12: 655–668.

29. Murray MG, Thompson WF. Rapid isolation of high molecular-weight plant DNA. Nucleic Acids Res. (1980); 8: 4321–4325.

30. Magoc T, Salzberg SL. FLASH: fast length adjustment of short reads to improve genome assemblies. Bioinformatics. (2011); 27: 2957–2963.

31. Bokulich NA, Subramanian S, Faith JJ, Gevers D, Gordon JI, Knight R, et al. Quality-filtering vastly improves diversity estimates from Illumina amplicon sequencing. Nat Methods. (2013); 10: 57–U11.

32. Edgar RC, Haas BJ, Clemente JC, Quince C, Knight R. UCHIME improves sensitivity and speed of chimera detection. Bioinformatics. (2011); 27: 2194–2200.

33. Haas BJ, Gevers D, Earl AM, Feldgarden M, Ward DV, Giannoukos G, et al. Chimeric 16S rRNA sequence formation and detection in Sanger and 454-pyrosequenced PCR amplicons. Genome Res. (2011); 21: 494–504.

34. Edgar RC. UPARSE: highly accurate OTU sequences from microbial amplicon reads. Nat Methods. (2013); 10: 996–998.

35. Quast C, Pruesse E, Yilmaz P, Gerken J, Schweer T, Yarza P, et al. The SILVA ribosomal RNA gene database project: improved data processing and web-based tools. Nucleic Acids Res. (2013); 41: D590–D596.

36. Chen YX, Chen YS, Shi CM, Huang ZB, Zhang Y, Li SK, et al. SOAPnuke: a MapReduce acceleration-supported software for integrated quality control and preprocessing of high-throughput sequencing data. Gigascience. (2017); 7.

37. Li DH, Liu CM, Luo RB, Sadakane K, Lam TW. MEGAHIT: an ultra-fast single-node solution for large and complex metagenomics assembly via succinct de Bruijn graph. Bioinformatics. (2015); 31: 1674–1676.

38. Wu YW, Simmons BA, Singer SW. MaxBin 2.0: an automated binning algorithm to recover genomes from multiple metagenomic datasets. Bioinformatics. (2016); 32: 605–607.

39. Kang DWD, Li F, Kirton E, Thomas A, Egan R, An H, et al. MetaBAT 2: an adaptive binning algorithm for robust and efficient genome reconstruction from metagenome assemblies. Peerj. (2019); 7.

40. Alneberg J, Bjarnason BS, de Bruijn I, Schirmer M, Quick J, Ijaz UZ, et al. Binning metagenomic contigs by coverage and composition. Nat Methods. (2014); 11: 1144–1146.

41. Uritskiy GV, DiRuggiero J, Taylor J. MetaWRAP-a flexible pipeline for genome-resolved metagenomic data analysis. Microbiome. (2018); 6.

42. Parks DH, Imelfort M, Skennerton CT, Hugenholtz P, Tyson GW. CheckM: assessing the quality of microbial genomes recovered from isolates, single cells, and metagenomes. Genome Res. (2015); 25: 1043–1055.

43. Dombrowski N, Teske AP, Baker BJ. Expansive microbial metabolic versatility and biodiversity in dynamic Guaymas Basin hydrothermal sediments. Nat Commun. (2018); 9.

44. Fardeau ML, Ollivier B, Patel BKC, Magot M, Thomas P, Rimbault A, et al. Thermotoga hypogea sp. nov., a xylanolytic, thermophilic bacterium from an oil-producing well. Int J Syst Bacteriol. (1997); 47: 1013–1019.

45. Buchan A, LeCleir GR, Gulvik CA, Gonzalez JM. Master recyclers: features and functions of bacteria associated with phytoplankton blooms. Nat Rev Microbiol. (2014); 12: 686–698.

46. Sekiguchi Y, Yamada T, Hanada S, Ohashi A, Harada H, Kamagata Y. Anaerolinea thermophila gen. nov., sp nov and Caldilinea aerophila gen. nov., sp nov., novel filamentous thermophiles that represent a previously uncultured lineage of the domain Bacteria at the subphylum level. Int J Syst Evol Micr. (2003); 53: 1843–1851.

47. Graham L, Orenstein JM. Processing tissue and cells for transmission electron microscopy in diagnostic pathology and research. Nat Protoc. (2007); 2: 2439–2450.

48. Zheng RK, Liu R, Shan YQ, Cai RN, Liu G, Sun CM. Characterization of the first cultured free-living representative of Candidatus Izimaplasma uncovers its unique biology. bioRxiv. (2020).

49. Loman NJ, Quinlan AR. Poretools: a toolkit for analyzing nanopore sequence data. Bioinformatics. (2014); 30: 3399–3401.

50. Koren S, Walenz BP, Berlin K, Miller JR, Bergman NH, Phillippy AM. Canu: scalable and accurate long-read assembly via adaptive k-mer weighting and repeat separation. Genome Res. (2017); 27: 722–736.

51. Richter M, Rossello-Mora R, Glockner FO, Peplies J. JSpeciesWS: a web server for prokaryotic species circumscription based on pairwise genome comparison. Bioinformatics. (2016); 32: 929–931.

52. Meier-Kolthoff JP, Auch AF, Klenk HP, Goker M. Genome sequence-based species delimitation with confidence intervals and improved distance functions. Bmc Bioinformatics. (2013); 14.

53. Wu DY, Jospin G, Eisen JA. Systematic identification of gene families for use as “markers” for phylogenetic and phylogeny-driven ecological studies of Bacteria and Archaea and their major subgroups. Plos One. (2013); 8.

54. Darling AE, Jospin G, Lowe E, Matsen FIV, Bik HM, Eisen JA. PhyloSift: phylogenetic analysis of genomes and metagenomes. Peerj. (2014); 2.

55. Trifinopoulos J, Nguyen LT, von Haeseler A, Minh BQ. W-IQ-TREE: a fast online phylogenetic tool for maximum likelihood analysis. Nucleic Acids Res. (2016); 44: W232–W235.

56. Letunic I, Bork P. Interactive Tree Of Life (iTOL): an online tool for phylogenetic tree display and annotation. Bioinformatics. (2007); 23: 127–128.

57. Letunic I, Bork P. Interactive tree of life (iTOL) v3: an online tool for the display and annotation of phylogenetic and other trees. Nucleic Acids Res. (2016); 44: W242–W245.

58. Yilmaz P, Parfrey LW, Yarza P, Gerken J, Pruesse E, Quast C, et al. The SILVA and “All-species Living Tree Project (LTP)” taxonomic frameworks. Nucleic Acids Res. (2014); 42: D643–D648.

59. Baker BJ, Appler KE, Gong X. New microbial biodiversity in marine sediments. Ann Rev Mar Sci. (2020).

60. Lewis WH, Tahon G, Geesink P, Sousa DZ, Ettema TJG. Innovations to culturing the uncultured microbial majority. Nat Rev Microbiol. (2020).

61. Zheng RK, Sun CM. Sphingosinithalassobacter tenebrarum sp. nov., isolated from a deep-sea cold seep. Int J Syst Evol Micr. (2020); 70: 5561–5566.

62. Henson MW, Lanclos VC, Faircloth BC, Thrash JC. Cultivation and genomics of the first freshwater SAR11 (LD12) isolate. ISME J. (2018); 12: 1846–1860.

63. Lloyd KG, Steen AD, Ladau J, Yin J, Crosby L. Phylogenetically novel uncultured microbial cells dominate Earth microbiomes. mSyetems. (2018); 3.

64. Yamada T, Sekiguchi Y. Cultivation of Uncultured Chloroflexi Subphyla: Significance and Ecophysiology of Formerly Uncultured Chloroflexi ‘Subphylum I’ with Natural and Biotechnological Relevance. Microbes Environ. (2009); 24: 205–216.

65. D’Hondt S, Rutherford S, Spivack AJ. Metabolic activity of subsurface life in deep-sea sediments. Science. (2002); 295: 2067–2070.

66. Xia Y, Lü C, Hou N, Xin Y, Liu J, Liu H, et al. Sulfide production and oxidation by heterotrophic bacteria under aerobic conditions. ISME J. (2017); 11: 2754.

