## Supplemental Figures and Tables for "Cultivation and characterization of a novel clade of deep-sea Chloroflexi: providing a glimpse of the phylum Chloroflexi involved in sulfur cycling"

### Supplementary methods

**Proteomic analysis**

**Sample processing protocol**

Strain ZRK33 was cultivated in the ORG medium supplemented without or with 200 mM $Na_2SO_4$ or 200 mM $Na_2S_2O_3$ for 8 d at 28 ℃ and then the cells were collected. These cells were sonicated three times on ice using a high intensity ultrasonic processor in lysis buffer (8 M urea, 1% Protease Inhibitor Cocktail). The remaining debris was removed by centrifugation at 12,000 *g* at 4 ℃ for 10 min. Finally, the supernatant was collected and the protein concentration was determined with a BCA kit (Solarbio, China) according to the manufacturer's instructions. For trypsin digestion, the protein solution was reduced with 5 mM dithiothreitol for 30 min at 56 ℃ and alkylated with 11 mM iodoacetamide for 15 min at room temperature in darkness. The 100 mm TEAB was added to the diluted protein sample in a solution with a urea concentration of less than 2 M. Finally, trypsin was added at a trypsin to protein mass ratio of 1:50 for the first digestion overnight, with 1:100 trypsin and protein. The mass was added for a second digestion for 4 h. Then the tryptic peptides were dissolved in 0.1% formic acid (solvent A) and directly loaded into a home-made reversed-phase analytical column (15-cm length, 75 μm inner diameter). The gradient increased from 6% to 23% in solvent B (0.1% formic acid in 98% acetonitrile) over 26 min, from 23% to 35% in 8 min and increased to 80% in 3 min, then maintain 80%

for the last 3 min, and all at a constant flow rate of 400 nL/min on an EASY-nLC 1000 UPLC system.

The peptides were coupled to UPLC in Q ExactiveTM Plus (Thermo, USA) via NSI source and tandem mass spectrometry (MS/MS). The applied electrospray voltage was 2.0 kV. The full scan has an m/z scan range of 350 to 1,800, and at 70,000 resolution, intact peptides were detected in the Orbitrap. MS/MS was then selected using the NCE set to 28 select peptides and fragments were detected in the Orbitrap at a resolution of 17,500. A data-related process that alternates between one MS scan followed by 20 MS/MS scans with 15.0 s dynamic exclusion. The automatic gain control (AGC) was set to 5E4. The fixed first mass was set as 100 m/z.

**Data processing protocol**

**(1) Database Search**

The resulting MS/MS data were processed using Maxquant search engine (v.1.5.2.8) [1]. Tandem mass spectra were searched against some databases (such as UniProt-GOA, InterPro, Kyoto Encyclopedia of Genes and Genomes (KEGG)) concatenated with reverse decoy database. Trypsin/P was specified as cleavage enzyme allowing up to 2 missing cleavages. The mass tolerance for precursor ions was set as 20 ppm in First search and 5 ppm in Main search, and the mass tolerance for fragment ions was set as 0.02 Da. Carbamidomethyl on Cys was specified as fixed modification and oxidation on Met was specified as variable modifications. FDR was adjusted to $< 1\%$ and minimum score for peptides was set $> 40$.

**(2) Enrichment of Gene Ontology analysis**

Proteins were classified by GO annotation into three categories: biological process, cellular compartment and molecular function. For each category, a two-tailed Fisher's exact test was employed to test the enrichment of the differentially expressed protein against all identified proteins. The GO with a corrected *P*-value < 0.05 is considered significant.

**(3) Enrichment of pathway analysis**

Encyclopedia of Genes and Genomes (KEGG) database was used to identify enriched pathways by a two-tailed Fisher's exact test to test the enrichment of the differentially expressed protein against all identified proteins [2]. The pathway with a corrected p-value < 0.05 was considered significant. These pathways were classified into hierarchical categories according to the KEGG website.

**(4) Enrichment of protein domain analysis**

For each category proteins, InterPro (a resource that provides functional analysis of protein sequences by classifying them into families and predicting the presence of domains and important sites) database was researched and a two-tailed Fisher's exact test was employed to test the enrichment of the differentially expressed protein against all identified proteins. Protein domains with a *P*-value < 0.05 were considered significant.

**(5) Enrichment-based Clustering**

For further hierarchical clustering based on different protein functional classification (such as: GO, Domain, Pathway, Complex). We first collated all the categories obtained after enrichment along with their *P* values, and then filtered for those categories which were at least enriched in one of the clusters with *P* value <0.05. This filtered *P* value matrix was transformed by the function $x = -\log_{10}$ (*P* value). Finally these x values were z-transformed for each functional category. These z scores were then clustered by one-way hierarchical clustering (Euclidean distance, average linkage clustering) in Genesis. Cluster membership was visualized by a heat map using the "heatmap.2" function from the "gplots" R-package.

### Supplementary results

**Description of Sulfochloroflexaceae fam. nov.**

Sulfochloroflexaceae (Sul.fo'ch.lo.ro.fle.xa.ce.ae. N.L. fem. n. *Sulfochloroflexus* type genus of the family; suff. -aceae, ending to denote a family; N.L. fem. pl. n. Sulfochloroflexaceae the family of the genus Sulfochloroflexus).

The description is the same as that for the genus *Sulfochloroflexus*. The type genus is *Sulfochloroflexus*.

**Description of Sulfochloroflexales ord. nov.**

Sulfochloroflexales (Sul.fo'ch.lo.ro.fle.xa.les. N.L. fem. n. Sulfochloroflexus type genus of the order; suff. -ales ending to denote an order; N.L. fem. pl. n. Sulfochloroflexales order of the genus *Sulfochloroflexus*).

The description is the same as that for the genus *Sulfochloroflexus*. The type genus is *Sulfochloroflexus*.

**Description of Sulfochloroflexia classis nov.**

Sulfochloroflexia (Sul.fo'ch.lo.ro.fle.xia. N.L. fem. n. Sulfochloroflexus type genus of the class; N.L. fem. pl. n. Sulfochloroflexia, the class of the order Sulfochloroflexales).

The class Sulfochloroflexia is defined on the basis of phylogenetic trees by comparative 16S rRNA gene, genome, RpoB and EF-tu sequences analysis from a wide variety of cultivated strains and environmental clones. The type order is Sulfochloroflexales.

### Supplementary figures

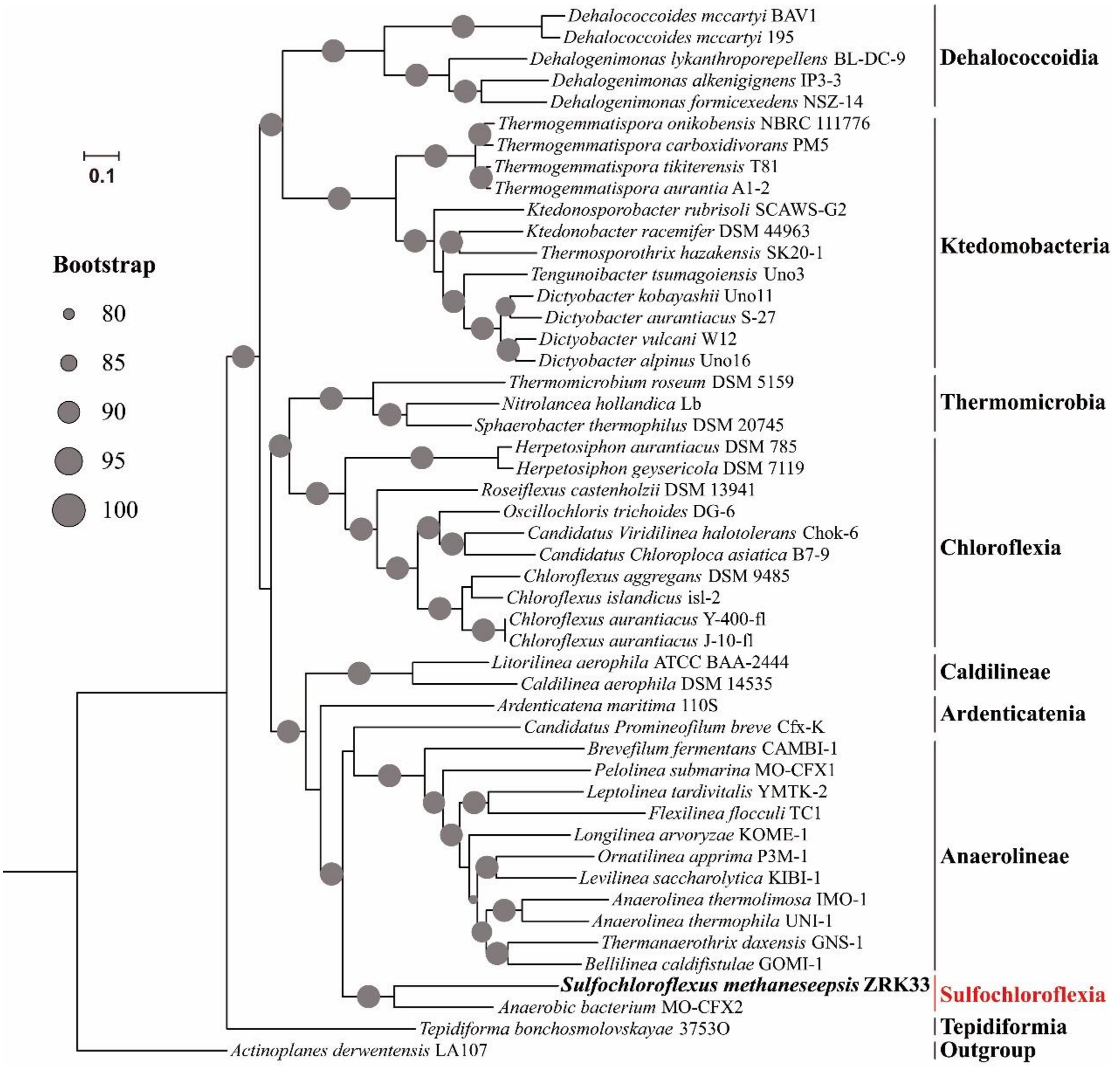


**Supplementary Figure S1.** Maximum likelihood phylogenetic tree of strain ZRK33 based on the genomes from all cultured Chloroflexi representatives using the concatenated alignment of 37 single-copy genes. *Actinoplanes derwentensis* LA107 was used as the outgroup. Nodes with greater than 80% bootstrap support are annotated with a black circle. Bar, 0.1 substitutions per nucleotide position.

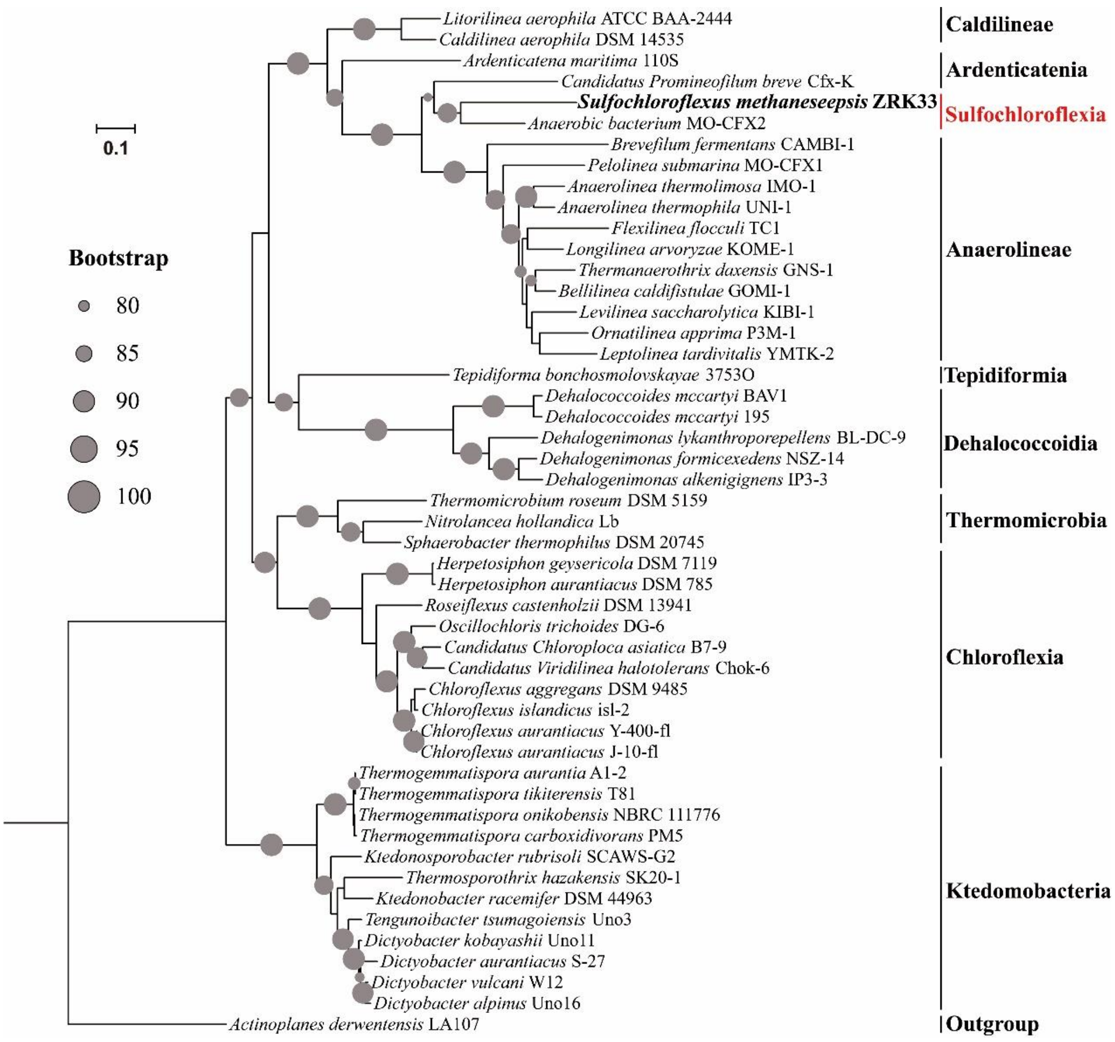


**Supplementary Figure S2.** Maximum likelihood phylogenetic tree of RpoB from genomes of strain ZRK33 and all cultured Chloroflexi representatives. *Actinoplanes derwentensis* LA107 was used as the outgroup. Nodes with greater than 80% bootstrap support are annotated with a black circle. Bar, 0.1 substitutions per nucleotide position.

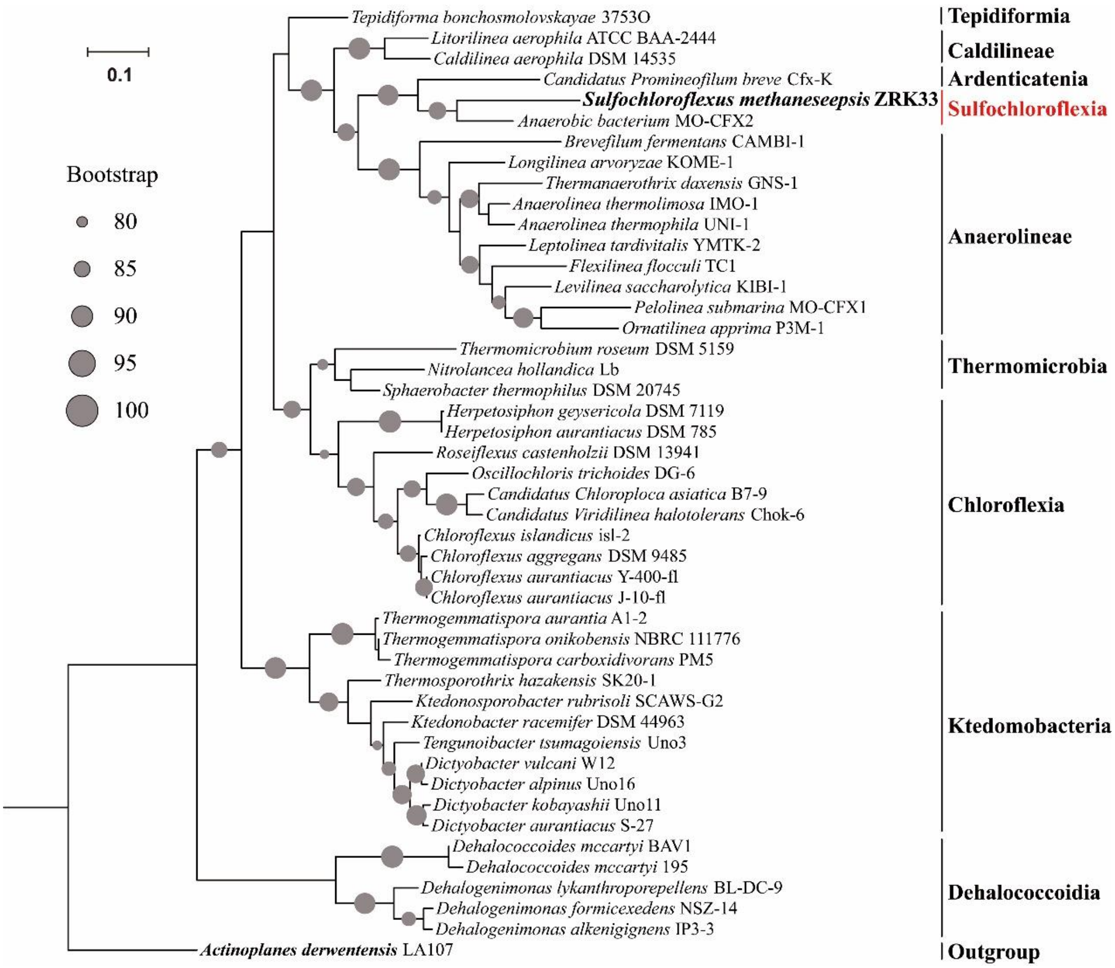


**Supplementary Fig. S3.** Maximum likelihood phylogenetic tree of elongation factor Tu (EF-Tu) from genomes of strain ZRK33 and all cultured Chloroflexi representatives. *Actinoplanes derwentensis* LA107 was used as the outgroup. Nodes with greater than 80% bootstrap support are annotated with a black circle. Bar, 0.1 substitutions per nucleotide position.

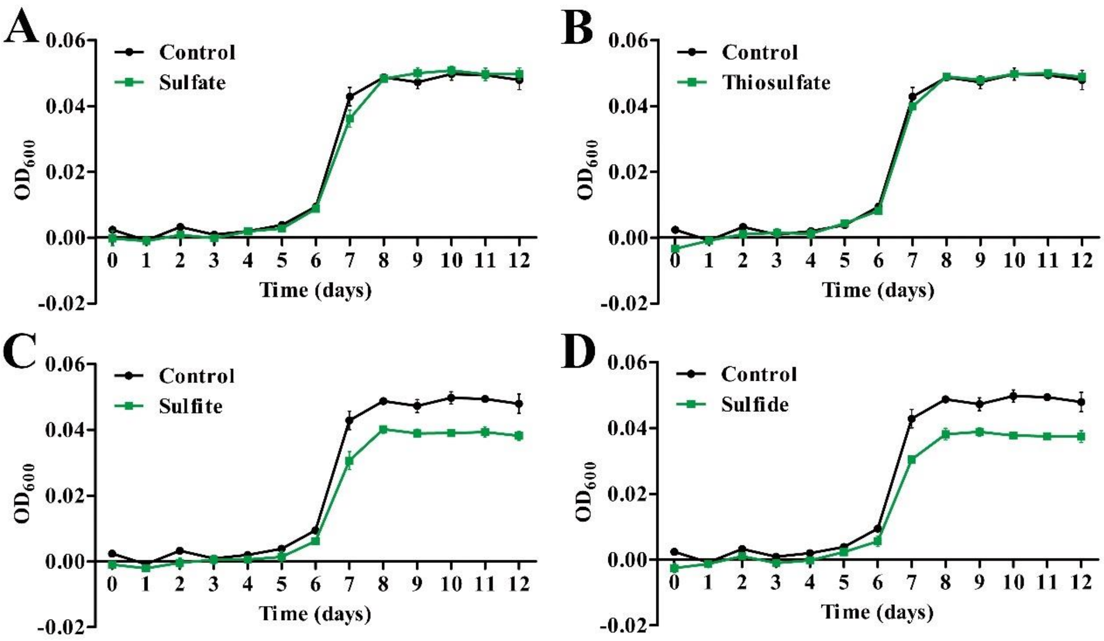


**Supplementary Fig. S4.** Growth assays of strain ZRK33 cultured in the medium supplemented with different sulfur-containing compounds. (A) Growth assays of strain ZRK33 in the medium supplemented without or with 20 mM $Na_2SO_4$. (B) Growth assays of strain ZRK33 in the medium supplemented without or with 20 mM $Na_2S_2O_3$. (C) Growth assays of strain ZRK33 in the medium supplemented without or with 1 mM $Na_2SO_3$. (D) Growth assays of strain ZRK33 in the medium supplemented without or with 1 mM $Na_2S$.

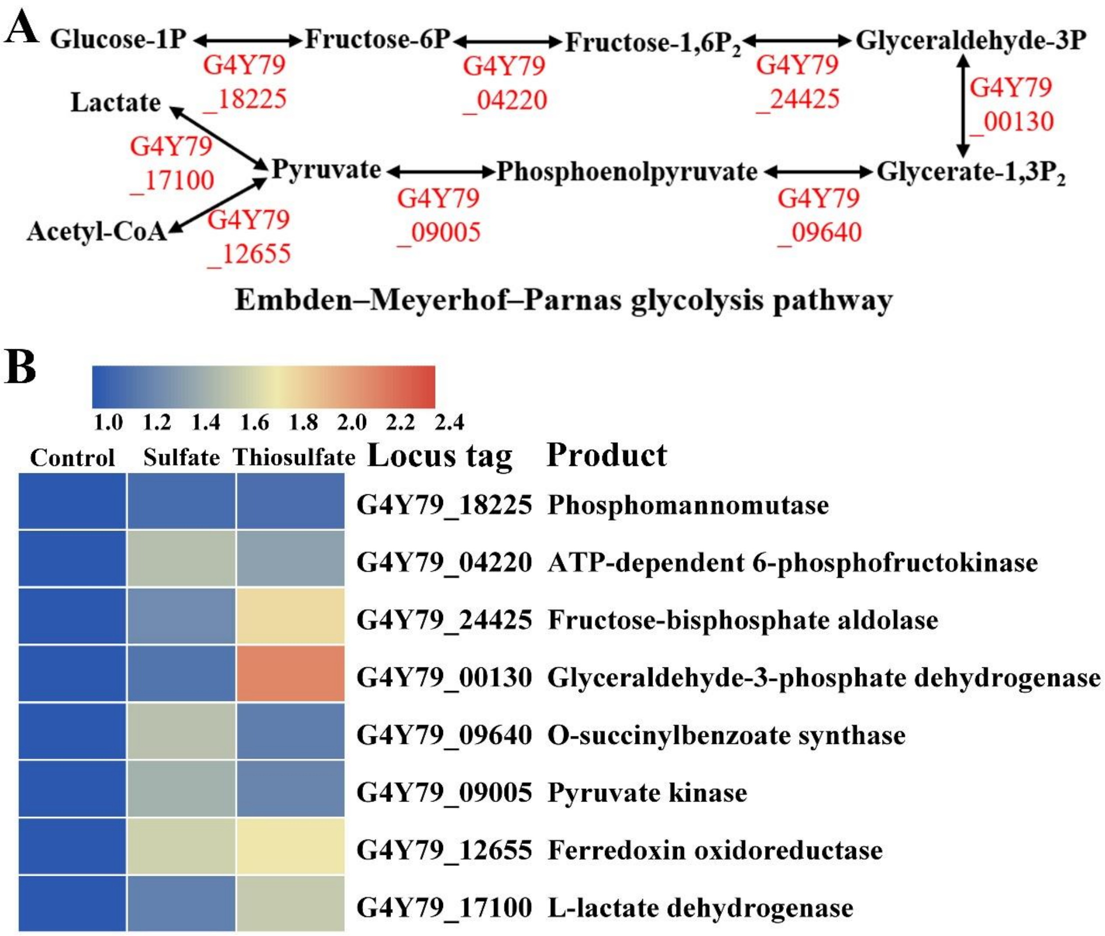


**Supplementary Fig. S5.** Proteomic analysis of expressions of genes associated with EMP glycolysis when strain ZRK33 was cultured in the medium supplemented with 200 mM sulfate or thiosulfate. (A) Diagrammatic scheme of EMP glycolysis pathway identified in the genome of strain ZRK33. The gene numbers showing in this scheme are extracted from the genome of strain ZRK33 (GenBank accession number CP051151) and they are the same with those shown in panel B. (B) Proteomics based heat map showing all up-regulated genes associated with EMP glycolysis pathway.

### Supplementary Tables

**Supplementary Table S1.** Marker genes used in phylogenetic analysis.

| ID | Protein |
|---|---|
| DNGNGWU00001 | ribosomal protein S2 rpsB |
| DNGNGWU00002 | ribosomal protein S10 rpsJ |
| DNGNGWU00003 | ribosomal protein L1 rplA |
| DNGNGWU00005 | translation initiation factor IF-2 |
| DNGNGWU00006 | metalloendopeptidase |
| DNGNGWU00007 | ribosomal protein L22 |
| DNGNGWU00009 | ribosomal protein L4/L1e rplD |
| DNGNGWU00010 | ribosomal protein L2 rplB |
| DNGNGWU00011 | ribosomal protein S9 rpsI |
| DNGNGWU00012 | ribosomal protein L3 rplC |
| DNGNGWU00013 | phenylalanyl-tRNA synthetase beta subunit |
| DNGNGWU00014 | ribosomal protein L14b/L23e rplN |
| DNGNGWU00015 | ribosomal protein S5 |
| DNGNGWU00016 | ribosomal protein S19 rpsS |
| DNGNGWU00017 | ribosomal protein S7 |
| DNGNGWU00018 | ribosomal protein L16/L10E rplP |
| DNGNGWU00019 | ribosomal protein S13 rpsM |
| DNGNGWU00020 | phenylalanyl-tRNA synthetase alpha subunit |
| DNGNGWU00021 | ribosomal protein L15 |
| DNGNGWU00022 | ribosomal protein L25/L23 |
| DNGNGWU00023 | ribosomal protein L6 rplF |
| DNGNGWU00024 | ribosomal protein L11 rplK |
| DNGNGWU00025 | ribosomal protein L5 rplE |
| DNGNGWU00026 | ribosomal protein S12/S23 |
| DNGNGWU00027 | ribosomal protein L29 |
| DNGNGWU00028 | ribosomal protein S3 rpsC |
| DNGNGWU00029 | ribosomal protein S11 rpsK |
| DNGNGWU00030 | ribosomal protein L10 |
| DNGNGWU00031 | ribosomal protein S8 |
| DNGNGWU00032 | tRNA pseudouridine synthase B |
| DNGNGWU00033 | ribosomal protein L18P/L5E |
| DNGNGWU00034 | ribosomal protein S15P/S13e |
| DNGNGWU00035 | Porphobilinogen deaminase |
| DNGNGWU00036 | ribosomal protein S17 |
| DNGNGWU00037 | ribosomal protein L13 rplM |
| DNGNGWU00039 | ribonuclease HII |
| DNGNGWU00040 | ribosomal protein L24 |

The DNGNGWU marker genes in phylosift refer to a suite of single-copy, protein-coding marker genes. All 37 DNGNGWU marker genes were concatenated to construct maximum likelihood phylogenetic tree.

**Supplementary Table S2.** Characteristics of strain ZRK33 and the other isolated Chloroflexi members. Strains: 1, strain ZRK33; 2, *Aggregatilinea lenta* MO-CFX2$^{T}$ [3]; 3, *Pelolinea submarina* MO-CFX1$^{T}$ [4]; 4, *Anaerolinea thermophila* UNI-1$^{T}$ [5]; 5, *Anaerolinea thermolimosa* IMO-1$^{T}$ [6]; 6, *Ornatilinea apprima* P3M-1$^{T}$ [7]. +, Positive; −, negative; NA, no data available.

| **Characteristic** | **1** | **2** | **3** | **4** | **5** | **6** |
|---|---|---|---|---|---|---|
| Cell morphology | Filaments | Filaments | Filaments | Filaments | Filaments | Filaments |
| Cell diameter (μm) | 0.3-0.5 | 0.5-0.6 | 0.13-0.15 | 0.2-0.3 | 0.3-0.4 | 0.3-0.7 |
| Temperature for growth (℃) | 28-32 | 20-37 | 10-37 | 50-60 | 42-55 | 20-50 |
| Optimum | 28 | 30 | 25-30 | 55 | 50 | 42-45 |
| pH for growth | 6.0-8.0 | 5.5-8.0 | 5.5-8.5 | 6.0-8.0 | 6.0-7.5 | 6.5-9.0 |
| Optimum | 7.0 | 6.5-7.0 | 7.0 | 7.0 | 7.0 | 7.5-8.0 |
| NaCl concentration for growth (%) | 0-5 | 0-3 | 0-5 | 0-5 | 0-2.5 | 0-2 |
| Draft (or complete) genome size (Mbp) | 5.6 | 6.2 | 3.5 | 3.5 | 4.2 | 4.4 |
| DNA G+C content (mol%) | 52.76 | 63.2 | 50.6 | 53.8 | 53.7 | 55.7 |
| Major cellular fatty acids | $C_{16:0}$,<br>$C_{15:0}$2-OH,<br>$C_{17:1}\omega$6c,<br>$C_{18:1}\omega$7c | $C_{16:0}$, $C_{18:0}$,<br>$C_{18:1}\omega$9c | $C_{18:1}\omega$9,<br>$C_{16:1}\omega$7,<br>i-$C_{17:0}$3-OH,<br>$C_{16:0}$ | $C_{16:0}$,<br>$C_{15:0}$, $C_{14:0}$ | ai-$C_{17:0}$,<br>i-$C_{15:0}$, $C_{16:0}$ | i-$C_{15:0}$,<br>ai-$C_{15:0}$,<br>$C_{14:0}$ |
| Doubling time | 4 h | 19 days | 1.5 days | 3 days | 2 days | 6 h |
| Substrates for growth: | | | | | | |
| Arabinose | + | − | + | + | + | − |
| Fructose | + | − | + | + | + | − |
| Glucose | + | − | + | + | + | + |
| Galactose | + | − | + | + | + | − |
| Mannose | + | − | − | + | + | NA |
| Ribose | + | − | + | + | + | NA |
| Xylose | − | − | + | + | + | + |
| Fumarate | + | + | − | − | − | − |
| Pyruvate | + | + | − | + | + | NA |
| Peptone | + | − | − | NA | + | − |
| Isolation source | Deep-sea cold seep sediments | Marine subsurface sediment | Marine subsurface sediment | Thermophilic anaerobic sludge | Thermophilic anaerobic sludge | Deep terrestrial hot aquifer |

**Supplementary Table S3.** Genomic features of strain ZRK33 with isolated Chloroflexi members.

| Feature | MO-CFX2 | MO-CFX1 | UNI-1 | IMO-1 | P3M-1 |
|---|---|---|---|---|---|
| ANIb (%) | 64.81 | 63.06 | 63.42 | 63.41 | 63.29 |
| ANIm (%) | 85.21 | 82.63 | 83.42 | 83.15 | 83.23 |
| Tetra | 0.48145 | 0.67572 | 0.64677 | 0.65234 | 0.65126 |
| GGDC (%) | 23.30 | 24.20 | 20.40 | 21.60 | 23.80 |

**Supplementary Table S4.** Assembly statistics and quality metrics of reconstructed genome bins of Chloroflexi used in this study.

| **Bin name** | **Taxonomy** | **Completeness (%)** | **Contamination (%)** | **GC (%)** | **N50 (bp)** | **Genome size (bp)** |
|---|---|---|---|---|---|---|
| zhu.bin.33 | Chloroflexi | 78.05 | 1.99 | 61.3 | 3481 | 2239783 |
| zhu.bin.3 | Chloroflexi | 56.84 | 1.98 | 0.494 | 4097 | 748332 |
| zhu.bin.7 | Chloroflexi | 59.57 | 1.98 | 0.446 | 5817 | 588020 |
| zhu.bin.9 | Chloroflexi | 66.38 | 1.925 | 0.506 | 2239 | 1167066 |
| zhu.bin.22 | Chloroflexi | 51.94 | 8.91 | 0.52 | 6548 | 1409231 |
| zhu.bin.44 | Chloroflexi | 66.88 | 0.99 | 0.528 | 6754 | 946329 |
| C1.bin.34 | Chloroflexi | 76.21 | 2.828 | 0.612 | 3548 | 2588152 |
| C1.bin.35 | Chloroflexi | 58.64 | 1.818 | 0.455 | 8245 | 1933721 |
| C2.bin.4 | Chloroflexi | 82.83 | 0 | 0.486 | 39431 | 941411 |
| C2.bin.6 | Chloroflexi | 70.92 | 0 | 0.495 | 7628 | 827319 |
| C2.bin.8 | Chloroflexi | 74.02 | 0.99 | 0.525 | 4817 | 757107 |
| C2.bin.9 | Chloroflexi | 80.36 | 1.98 | 0.548 | 6764 | 1051572 |
| C2.bin.12 | Chloroflexi | 54.49 | 2.727 | 0.523 | 3759 | 1643107 |
| C2.bin.17 | Chloroflexi | 65.4 | 4.158 | 0.542 | 4882 | 621181 |
| C2.bin.33 | Chloroflexi | 63.82 | 1.386 | 0.609 | 3494 | 1094429 |
| C2.bin.34 | Chloroflexi | 62.68 | 2.727 | 0.479 | 4652 | 2209326 |
| C2.bin.38 | Chloroflexi | 72.49 | 4.022 | 0.619 | 3598 | 2727830 |
| C2.bin.45 | Chloroflexi | 87.29 | 1.485 | 0.537 | 9527 | 1647588 |
| C2.bin.48 | Chloroflexi | 61.22 | 8.25 | 0.452 | 5264 | 973927 |
| C4.bin.19 | Chloroflexi | 67.43 | 0.565 | 0.644 | 3026 | 1749317 |
| H1.bin.7 | Chloroflexi | 73.68 | 0 | 0.545 | 4996 | 1405141 |
| H1.bin.32 | Chloroflexi | 71.94 | 7.727 | 0.563 | 3845 | 2766581 |
| H2.bin.45 | Chloroflexi | 76.73 | 4.378 | 0.579 | 3649 | 942250 |
| H2.bin.80 | Chloroflexi | 86.57 | 0.925 | 0.663 | 14655 | 3235763 |
| H2.bin.87 | Chloroflexi | 59.82 | 1.485 | 0.603 | 3209 | 1355795 |
| H2.bin.116 | Chloroflexi | 92.73 | 0.99 | 0.543 | 27621 | 1910274 |
| H2.bin.125 | Chloroflexi | 70.13 | 0.99 | 0.477 | 5832 | 1871708 |

#### References

1. Cox J, Mann M. MaxQuant enables high peptide identification rates, individualized p.p.b.-range mass accuracies and proteome-wide protein quantification. Nat Biotechnol. (2008); 26: 1367-1372.
2. Kanehisa M, Sato Y, Kawashima M, Furumichi M, Tanabe M. KEGG as a reference resource for gene and protein annotation. Nucleic Acids Res. (2016); 44: D457-D462.
3. Nakahara N, Nobu MK, Takaki Y, Miyazaki M, Tasumi E, Sakai S, et al. *Aggregatilinea lenta* gen. nov., sp. nov., a slow-growing, facultatively anaerobic bacterium isolated from subseafloor sediment, and proposal of the new order Aggregatilineales ord. nov. within the class Anaerolineae of the phylum Chloroflexi. Int J Syst Evol Micr. (2019); 69: 1185-1194.
4. Imachi H, Sakai S, Lipp JS, Miyazaki M, Saito Y, Yamanaka Y, et al. *Pelolinea submarina* gen. nov., sp nov., an anaerobic, filamentous bacterium of the phylum Chloroflexi isolated from subseafloor sediment. Int J Syst Evol Micr. (2014); 64: 812-818.
5. Sekiguchi Y, Yamada T, Hanada S, Ohashi A, Harada H, Kamagata Y. *Anaerolinea thermophila* gen. nov., sp nov and *Caldilinea aerophila* gen. nov., sp nov., novel filamentous thermophiles that represent a previously uncultured lineage of the domain Bacteria at the subphylum level. Int J Syst Evol Micr. (2003); 53: 1843-1851.
6. Yamada T, Sekiguchi Y, Hanada S, Imachi H, Ohashi A, Harada H, et al. *Anaerolinea thermolimosa* sp nov., *Levilinea saccharolytica* gen. nov., sp nov and *Leptolinea tardivitalis* gen. nov., so. nov., novel filamentous anaerobes, and description of the new classes anaerolineae classis nov and Caldilineae classis nov in the bacterial phylum Chloroflexi. Int J Syst Evol Micr. (2006); 56: 1331-1340.
7. Podosokorskaya OA, Bonch-Osmolovskaya EA, Novikov AA, Kolganova TV, Kublanov IV. *Ornatilinea apprima* gen. nov., sp nov., a cellulolytic representative of the class Anaerolineae. Int J Syst Evol Micr. (2013); 63: 86-92.
